# Temperature and day length drive local adaptation in the Patagonian foundation tree species *Nothofagus pumilio*

**DOI:** 10.1101/2023.04.28.538677

**Authors:** Jill Sekely, Paula Marchelli, Verónica Arana, Benjamin Dauphin, María Gabriela Mattera, Mario Pastorino, Ivan Scotti, Carolina Soliani, Katrin Heer, Lars Opgenoorth

## Abstract

Climate change alters relationships among environmental conditions and thus has the potential to change the selection pressures acting on adaptive gene variants. Using a landscape genomic approach, we show that the southern beech species *Nothofagus pumilio* has notable genetic adaptations to climate along its 2000-kilometer-long range in the Andes. We screened 47,336 SNP loci in 1,632 contigs and found that high-latitude sampling sites have lower genetic diversity, likely due to greater impact of glacial oscillations at high latitudes. Using four genome scan methods, we identified 457 outlier SNPs that are either strongly differentiated among subpopulations or associated with environmental covariates related to temperature, day length, and precipitation. Temperature and day length parameters were associated with notably more outliers than precipitation (n = 133, 113, and 61 outliers, respectively), and almost half of all annotated outliers were related to stress response (n=38, 21%) or catabolism-metabolism (n=43, 24%). Our findings suggest that *Nothofagus pumilio* is an ideal Andean model of genetic adaptation to climate change because it is locally adapted to extant climate conditions, and shifting patterns among environmental parameters may be detrimental to its future survival and adaptation potential.

## Introduction

Contemporary climate change is expected to have acute impacts on forests, including shifts in species ranges, tree growth rates, and phenology (Brondizio, Settele, Díaz, & Ngo, 2019). Tree populations typically have high levels of standing genetic diversity due to their widespread distributions across diverse habitats and large effective population sizes, and therefore they may have high local adaptation potential even when faced with rapidly changing conditions (Kremer et al., 2012; Savolainen, Pyhäjärvi, & Knürr, 2007). However, the extraordinary challenge of contemporary climate change is that it will likely create novel combinations of precipitation, temperature, and photoperiod that neither occur within the current range nor have occurred for millions of years (Burke et al., 2018; Williams & Jackson, 2007). By decoupling current relationships among environmental conditions, no-analog conditions could impose unique selection pressures that will challenge tree populations’ ability to survive.

While photoperiod is unaffected by climate change, temperature and precipitation patterns will shift across regions (Barros et al., 2015; Williams & Jackson, 2007). The consequences are myriad. In extratropical species, phenology is mediated by a combination of photoperiod and temperature cues (Howe, Hackett, Furnier, & Klevorn, 1995; Singh, Svystun, AlDahmash, Jönsson, & Bhalerao, 2017). Climate shifts also have implications for drought. Drought is a direct consequence of water availability, but its severity is influenced by temperature, since high temperatures can increase evapotranspiration rates and drought stress during the growing season (Vicente-Serrano, Beguería, & López-Moreno, 2010). Furthermore, pests and pathogen species including insects, bacteria, and fungi may likewise experience range or phenology shifts due to climate change. Therefore, no-analog climate combinations will likely affect many genes and traits related to phenology (Hänninen & Tanino, 2011), extreme temperature and drought response (e.g. Niinemets, 2010), and immune response (Haynes, Liebhold, Lefcheck, Morin, & Wang, 2022). Selection upon these genes leaves signatures of adaptation along the genome, and searching for signatures among putative adaptive loci can provide critical information about how tree populations might respond to climate change. An initial step is to establish whether adaptation to environmental clines is currently observed.

The Patagonia region of the southern Andes mountain range presents an ideal study location due to its north-south orientation and two geographically orthogonal environmental gradients. The first is a north-south gradient of day-length and temperature that is driven by latitude, and the second is a west-east precipitation gradient driven by prevailing winds and a montane rain shadow. Patagonia is strongly affected both by glacial oscillations and the El Niño-Southern Oscillation (Morales et al., 2020). Climate change has already intensified the acute drought stress following La Niña events (Cai et al., 2015). The most widespread native tree species in this region is the southern beech “lenga” (*Nothofagus pumilio* ([Poepp. & Endl.] Krasser)), a cold-tolerant deciduous tree that inhabits a nearly continuous range more than 2,000 kilometers long. Its range encompasses a diverse climate space, from 5,000 mm of annual precipitation in the west to just 200 mm in the east (Veblen, Donoso, Kitzberger, & Rebertus, 1996; Fig 1). Little is known about adaptation patterns at the fine geographical scale in this non-model montane species. Previous studies have examined the neutral genetic diversity and phenotypic plasticity using neutral markers(Arana et al., 2016; Mathiasen & Premoli, 2013, 2016; A C Premoli, 2003). However, adaptive variation along environmental clines using high-throughput SNPs has not yet been assessed.

**Figure 1.**
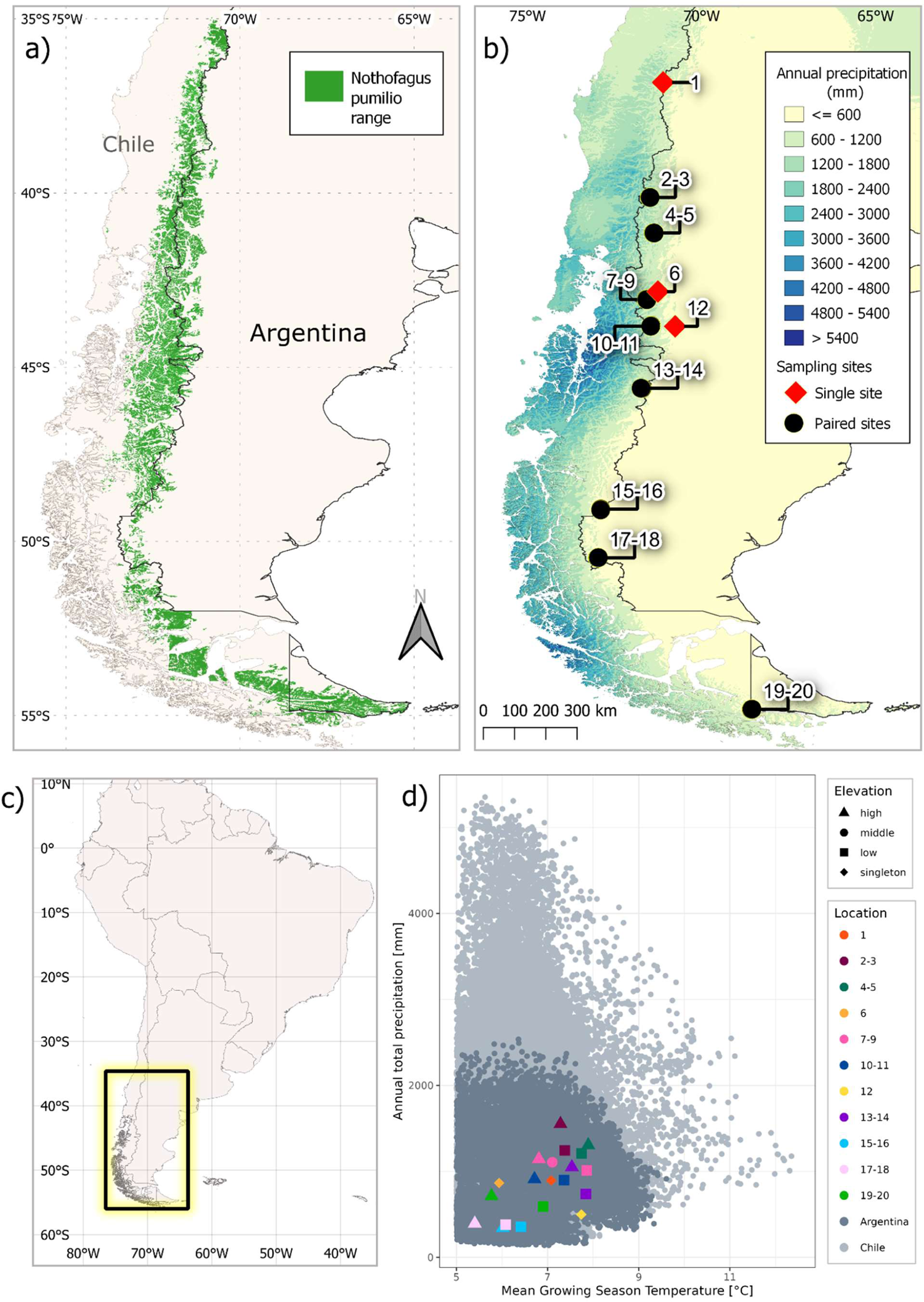
Characterisation of the *Nothofagus pumilio* distribution and study area. a) species distribution map, b) example of a bioclim layer from CHELSA (total annual precipitation) with sampling sites, c) overview map, d) climate space inhabited by *N. pumilio* in terms of mean temperature of all growing season days (℃) based on TREELIM and mean annual total precipitation (mm). Both values are the mean across years 1979-2013. Values are shown for the full distribution range in Argentina and Chile and sampling sites are grouped by color, with local elevation classes within sites differentiated by shape. Mean growing season temperature was truncated at 5℃ to compensate for low spatial resolution of CHELSA data, which showed erroneous low-temperature artifacts at high elevations due to sharp mountain slopes.

The objective of this study was to characterize extant local adaptation in *N. pumilio*. We searched for signatures of local adaptation within candidate genes in situ using a landscape genomics approach to assess how evolutionary processes and environmental variation have shaped genetic variation (Capblancq & Forester, 2021; Rellstab, Gugerli, Eckert, Hancock, & Holderegger, 2015). Local adaptation depends on a fine balance among many factors whose individual effects can be difficult to differentiate. Thus, choosing an appropriate sampling design and analysis methods is crucial for improving study power to detect signatures (Lotterhos & Whitlock, 2015; Meirmans, 2015). We used a paired-site sampling design, which aims to disentangle environmental effects from neutral population structure by maximizing the climatic distance between pairs of sampling sites while minimizing the neutral genetic divergence (Lotterhos & Whitlock, 2015; Scotti et al., 2023). We distributed sampling site pairs along the two orthogonal gradients on the eastern side of the Andes, and assessed variation using univariate and multivariate genome scan methods that incorporate population structure. We hypothesize that the two gradients have exerted strong selection pressure on *N. pumilio* and have resulted in signatures of local adaptation in candidate genes. We address these questions by quantifying the strength of correlations between environmental covariate predictors and genetic SNP responses. We predicted that (i) allele frequencies in candidate genes that are linked to growth and stress response will correlate with temperature and photoperiod clines and (ii) allele frequencies in candidate genes linked to drought response will correlate with precipitation, albeit to a weaker degree given the narrower precipitation gradient covered.

## Materials and Methods

### Study species

*Nothofagus pumilio*, common name “lenga,” is a deciduous tree species native to the southernmost temperate forest of the Andes mountains. It is a wind-pollinated and strictly outcrossing species that grows between latitudes 35° to 56 °S (Veblen et al., 1996). A member of the Fagaceae family, its closest relatives in the Northern Hemisphere are *Fagus* species (Vento & Agraín, 2018). It is cold-tolerant and often forms monospecific stands up to the montane tree line. North of 41°S, lenga grows in the subalpine zone, but it also grows at sea level in the southernmost (i.e. poleward) parts of its range. *Nothofagus pumilio* is an important local forestry species, although much of the local timber industry has historically focused on introduced genera from the Northern Hemisphere such as *Pinus* and *Eucalyptus* (Gea-Izquierdo, Pastur, Cellini, & Lencinas, 2004). It is a non-model species without a reference genome and a de novo transcriptome was recently assembled (Estravis-Barcala et al., 2021).

### Sampling design

To disentangle neutral and adaptive genetic variation, we used a paired-site study design after Lotterhos and Whitlock (2015). Sites within a pair are geographically close enough to share a demographic history but are distant enough that they experience different environmental selection pressures. According to Lotterhos and Whitlock (2015), this sampling design has greater power to detect signatures of local adaptation compared to transect or random sampling designs, particularly when combined with genome scan methods based on latent factor mixed models (LFMM) and Bayesian methods (see genome scan methods below). We selected eight localities that were distributed along the species’ full latitudinal range on the eastern slope of the Andes (Fig 1), and each locality contained two (or, in one locality, three) sampling sites. Linear distance among paired sites within a locality was always less than three kilometers, and elevation difference among the sites’ centroids was between 150 - 320 meters to capture an approximate 1-2 °C difference in mean annual temperature due to lapse rate (Whiteman, 2000). The high-elevation sites were located below the alpine treeline to avoid sampling trees with shrub-like krummholz formation (Table 1). In addition, we sampled three singleton localities in marginal habitats (i.e. located at the edge of the species distribution). Each singleton locality has one sampling site (Epulaufquen (site 1), La Hoya (6), and Jose de San Martin (12)). The latter two sites are located at approximately the same latitude as one of the paired localities, in an attempt to capture a wider portion of the east-west precipitation gradient.

**Table 1.**
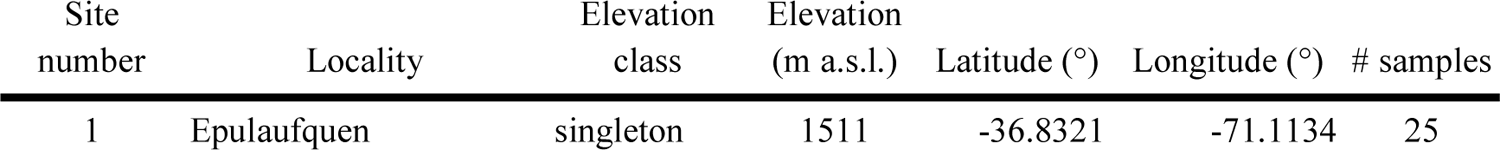
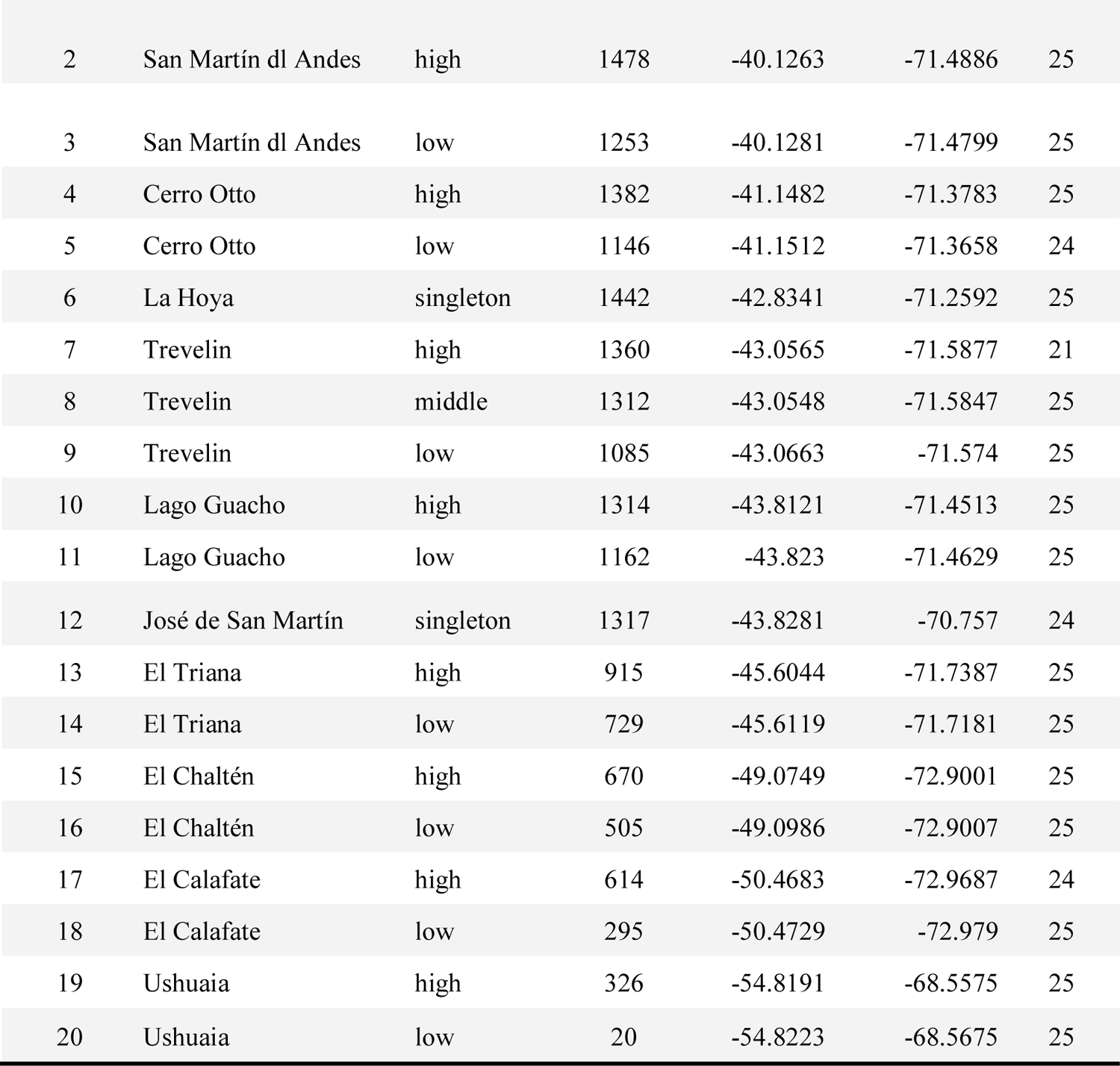
Characteristics of the *Nothofagus pumilio* sampling sites, ordered from North (site number 1) to South (20).

| Site number | Locality | Elevation class | Elevation (m a.s.l.) | Latitude (°) | Longitude (°) | # samples |
| --- | --- | --- | --- | --- | --- | --- |
| 1 | Epulaufquen | singleton | 1511 | -36.8321 | -71.1134 | 25 |
| 2 | San Martín dl Andes | high | 1478 | -40.1263 | -71.4886 | 25 |
| 3 | San Martín dl Andes | low | 1253 | -40.1281 | -71.4799 | 25 |
| 4 | Cerro Otto | high | 1382 | -41.1482 | -71.3783 | 25 |
| 5 | Cerro Otto | low | 1146 | -41.1512 | -71.3658 | 24 |
| 6 | La Hoya | singleton | 1442 | -42.8341 | -71.2592 | 25 |
| 7 | Trevelin | high | 1360 | -43.0565 | -71.5877 | 21 |
| 8 | Trevelin | middle | 1312 | -43.0548 | -71.5847 | 25 |
| 9 | Trevelin | low | 1085 | -43.0663 | -71.574 | 25 |
| 10 | Lago Guacho | high | 1314 | -43.8121 | -71.4513 | 25 |
| 11 | Lago Guacho | low | 1162 | -43.823 | -71.4629 | 25 |
| 12 | José de San Martín | singleton | 1317 | -43.8281 | -70.757 | 24 |
| 13 | El Triana | high | 915 | -45.6044 | -71.7387 | 25 |
| 14 | El Triana | low | 729 | -45.6119 | -71.7181 | 25 |
| 15 | El Chaltén | high | 670 | -49.0749 | -72.9001 | 25 |
| 16 | El Chaltén | low | 505 | -49.0986 | -72.9007 | 25 |
| 17 | El Calafate | high | 614 | -50.4683 | -72.9687 | 24 |
| 18 | El Calafate | low | 295 | -50.4729 | -72.979 | 25 |
| 19 | Ushuaia | high | 326 | -54.8191 | -68.5575 | 25 |
| 20 | Ushuaia | low | 20 | -54.8223 | -68.5675 | 25 |

We sampled between 21 and 25 adult trees per site for a total of 496 individuals. Selected trees were dominant or co-dominant and at least 50 years old, as confirmed by annual tree rings (Sekely et al, manuscript in progress). Intertree distances were at least 30 meters to reduce the chance of sampling directly related individuals. Geographic coordinates for each tree were recorded with a handheld GPS device (Garmin model GPSMAP 64st). We collected fresh leaf buds for DNA extraction and stored them at −80°C. Immediately before extraction, buds were manually descaled, flash-frozen with liquid nitrogen, and ground with mortar and pestle. Samples were randomly assigned to extraction batches. Total genomic DNA was extracted from 0.1 g of plant material using the CTAB protocol by Doyle (1990) with minor modifications, since *N. pumilio* leaf buds have high levels of polysaccharides and polyphenols that can impact the quality and quantity of extracted DNA. Therefore we added 1% soluble Polyvinylpyrrolidone (PVP) and Dithiothreitol (DTT) to the lysis buffer (Porebski, Bailey, & Baum, 1997). Extracted DNA quantity was measured with a QUBIT 1.0 Fluorometer (Invitrogen, Carlsbad, CA) and the quality was spot-checked with Nanodrop™ 2000 (ThermoFisher Scientific, catalog ND-2000). Extracted DNA samples were randomized among plates for downstream sequencing.

### Environmental data and covariate choice

Empirical climate data is limited for the Andes region, so environmental covariates were extracted from the public repository climate dataset CHELSA v.1.2 (Karger et al., 2017). CHELSA incorporates empirical climate data from 1979-2013, from which further bioclim variables were derived and extrapolated across the globe at a resolution of 30 arc sec (∼1 km^2^). We chose this dataset since it has been shown to represent more accurate orographic conditions than WorldClim (e.g. Bobrowski, Weidinger, & Schickhoff, 2021). We extracted tree-level data from climate layers with R::raster package (v. 3.6.14) using the extract() command for individual tree GPS locations and the “bilinear” option, which interpolates values from the four nearest raster cells to approximate finer-scale climate parameters (Hijmans, 2023).

Genome scans are sensitive to collinearity, so covariate pruning prior to analysis is a critical step (Dormann et al., 2013; Rellstab et al., 2015). From the CHELSA dataset we first selected a short list of variables related to temperature and drought stress (Fig S1). We calculated pairwise Pearson correlation values among these covariates, using the R package psych (v 2.2.9, Revelle, 2022), and retained only the most relevant variables that had a value less than |0.8| (Fig S2). Ultimately, we selected two CHELSA temperature parameters (number of frost days and isothermality (bioclim 3)), one precipitation parameter (annual precipitation amount (bioclim 12)) and one temperature-limited precipitation parameter (precipitation during growing season (gsp_9)). Finally, since we also investigated circadian clock candidate genes, we calculated average day length in the midsummer month of January (dl.Jan) using the R package geosphere (v 1.5.18, Hijmans, 2022). All environmental variables were scaled prior to association analyses.

### Probe design

Trees were genotyped with targeted sequencing (i.e. exome capture), for which we assembled a starting set of candidate genes. A recent study investigated 811 candidate gene orthogroups in seven European tree species that are closely- or distantly-related to *N. pumilio*, including Fagaecea members (Milesi et al., 2023 in review; Opgenoorth et al., 2021). Candidate genes were pertinent to environmental variables and were therefore selected from Gene Ontology (GO) and KEGG gene regulation networks related to cold, heat, drought, and immune response (Ashburner et al., 2000; Carbon et al., 2009). These orthogroups are represented by 1,789 candidate genes in *Arabidopsis thaliana*. We additionally investigated 415 species-specific *N. pumilio* candidate genes, including those that were differentially expressed in a recent heat stress transcriptomic study (Estravis-Barcala et al., 2021) or are affiliated with wood growth or circadian clock rhythms (Estravis-Barcala et al., 2020).

We used BLASTn (Altschul, Gish, Miller, Myers, & Lipman, 1990) to align the de novo *N. pumilio* transcriptome with the 2,204 candidate genes, retained the best transcriptome hit per gene by applying an e-value threshold of 10^−5^, and finally selected the best sequence hit per gene. This resulted in 1,467 contig hits that covered 2.58 Mb. To reach the probe design target size of 3 Mb, we complemented the list with the longest available contigs from the *N. pumilio* transcriptome. The final probe design encompassed 1,913 contigs, each containing between 200 - 11,873 nucleotides.

### Library preparation and target sequencing

Libraries were prepared with SeqCap EZ-HyperPlus (Roche Sequencing Solutions). Library size was analyzed with Bioanalyzer High Sensitivity DNA assay (Agilent technologies), library quantity was analyzed with Qubit 2.0 Fluorometer, and libraries were sequenced on NovaSeq 6000 (Illumina) in pair-end mode with 150 cycles per read. A total of 3.1 trillion reads were produced. Base-calling and demultiplexing were completed with Illumina bcl2fastq v.2.20. Reads were trimmed using ERNE v1.4.6 (Del Fabbro, Scalabrin, Morgante, & Giorgi, 2013) and cutadapt (Martin, 2011) then mapped onto the transcriptome with BWA-MEM v0.7.17 (Li & Durbin, 2009). Variants were called using gatk-4.0 (Poplin et al., 2017), first with HaplotypeCaller and then joint genotyping was performed using GenotypeGVCFs (DePristo et al., 2011). Variants were selected with GATK SelectVariants and coarsely quality-filtered in VariantFiltration. Default parameters were used for each step.

### Variant quality filtering

Variants were quality-filtered using general best practice thresholds in vcftools v 0.1.16 (Danecek et al 2011). These thresholds were minimum read depth per locus > 8 (command: --minDP 8), minimum quality > 20 (--minGQ 20), and maximum missing data per locus 20% (--max-missing 0.8) (Carson et al., 2014). We calculated genotype missingness per individual (--imiss). GATK in docker mode was used to remove all newly-created monomorphic loci and all individuals with > 50% missing data (n = 3 individuals). Multiallelic loci were removed, leaving only biallelic loci (--min-alleles 2 and --max-alleles 2). After these initial quality filtering steps, our dataset contained 116,136 SNPs in 1,783 contigs.

Paralogous loci were identified with the HDplot method and its accompanying R script (McKinney, Waples, Seeb, & Seeb, 2017), then pruned based on author recommendations (H > 0.6 and/or D > |20|). We used R version 4.2.2 for all analyses (R Core Team, 2022). We pruned the dataset of loci in linkage disequilibrium using plink v1.9 (Chang et al., 2015) to retain the allele with the greater minor allele frequency. We used the following settings: window size 50, stepwise progression 10, and r^2^ threshold 0.5 (plink v1.9 --indep-pairwise 50 10 0.5). This pipeline created our “main dataset,” which contained 47,336 SNPs (in 1,632 contigs). As a subsequent step, we applied a minor allele frequency filter of 5% to create a “maf-filtered dataset” that contained 9,601 SNPs (1,437 contigs).

### Descriptive genetic diversity statistics and population structure

We ran all genetic diversity and population structure analyses using the main dataset, with the exception of nucleotide diversity. Pairwise *F*_ST_ statistics were calculated in vcftools for every possible pair of sampling sites (n=190) with the weighted θ correction (Weir & Cockerham, 1984). The rarefied count of private alleles was calculated with the R::poppr package (v 2.9.3, Kamvar, Tabima, & Grünwald, 2014). We calculated heterozygosity and *F*_IS_ using hierfstat (v 0.5.11, Goudet & Jombart, 2022). Nucleotide diversity was calculated with pixy (v 1.2.7; Korunes & Samuk, 2021), which includes invariant sites to calculate unbiased values. Therefore our input dataset contained the main dataset plus every called invariant site, which were quality-filtered using the same thresholds. Per pixy user guidelines, we aggregated values within a sampling site by summing raw count differences and dividing by summed comparisons. Pearson correlation coefficients between genetic diversity statistics and latitude were calculated using ggpubr package and stat_cor command (v 0.5.0, Kassambara, 2022). Population structure was analyzed using the ADMIXTURE software (Alexander, Novembre, & Lange, 2009). We assessed every K value from 1 to 20, to represent the 20 sampling sites. Singletons can confound model-based inference of population structure such as ADMIXTURE (Linck & Battey, 2019), so they were removed prior to analysis.

### Genome scan methods

To determine whether SNPs are under selection, we applied genome scan methods to the maf-filtered dataset. Genome scans compare genetic variation of SNP loci across the targeted genome areas and identify over-differentiated loci, hereafter called outlier SNPs. There is an ever-growing list of genome scan tools and algorithms (see Bourgeois & Warren, 2021), each of which has its own benefits and pitfalls (e.g. Rellstab et al., 2015; Waldvogel, Schreiber, Pfenninger, & Feldmeyer, 2020). Common practice is to analyze a SNP dataset with multiple methods and inspect overlap among their results, since this provides stronger evidence that a SNP is a true-positive outlier (de Villemereuil, Frichot, Bazin, François, & Gaggiotti, 2014; Waldvogel et al., 2020).

Genome scans search for loci that are strongly differentiated among genetic clusters (e.g. subpopulations) and/or strongly associated with environmental gradients (Savolainen, Lascoux, & Merilä, 2013). We use both methods and classify them respectively as “population differentiation” (sensu Beaumont & Nichols, 1996) and “genotype-environment association” (sensu Hedrick, Ginevan, & Ewing, 1976). Population differentiation (PD) tests are advantageous because they require no prior knowledge about environmental selection pressures and therefore are less susceptible to errors related to missing environmental data or suboptimal choice of climatic variables. We used pcadapt, which is a multiple linear regression method that identifies outlier loci via correlation to genetic structure ordination axes (Duforet-Frebourg, Bazin, & Blum, 2014) and is implemented in the R package pcadapt (v 4.3.3, Privé, Luu, Vilhjálmsson, & Blum, 2020). The number of principal components (K = 3) was chosen based on the lowest genomic inflation factor (lambda = 1.35) and the deflation of explained variance of the first three principal components. On the other hand, genotype-environment associations (GEA) can provide evidence about which environmental variables are associated with adaptive differentiation. The null hypothesis in a GEA is that there is no correlation between allele frequencies and environmental covariates (Manel et al., 2010). GEA methods have greater power than PD to detect weakly selected loci, which may only show small allele frequency shifts but are crucial for adaptation (e.g. De La Torre, Wilhite, & Neale, 2019). We used three GEA methods that assess SNP frequency variations and environmental covariates in different combinations of univariate and multivariate approaches.

The Bayesian hierarchical model BayPass (v. 2.31, Gautier, 2015) is a univariate method for both genetic and environmental components. It computes X_T_X values, which are analogous to the SNP-specific *F*_ST_ that is calculated by PD methods, and Bayes Factor (BF), which measures the strength of correlation between an individual SNP and an individual environmental covariate. BayPass is a stochastic algorithm, so we ran three iterations with the core model (i.e. without environmental covariates) to calculate the population covariance matrix and X_T_X values, then calculated median values of each statistic. Next we ran three iterations of the auxiliary covariate model using the median covariate matrix to calculate Bayes Factor values and again retained median values. An important distinction is that BayPass requires allele frequencies to be pooled by sampling site, thus treating each of the 20 sites as its own population, unlike the other GEA methods.

The two additional GEA methods assess multivariate environmental parameters to account for interaction among environmental factors across the Patagonian landscape. The first is latent factor mixed models (LFMMs), which search for correlations between an individual SNP and multivariate environmental predictors while simultaneously correcting for hidden (i.e. latent) factors (Frichot, Schoville, Bouchard, & François, 2013). Latent factors can include unobserved demographic patterns or unmeasured environmental variables. This method is demonstrably effective in continuous ranges and has low false positive rates even if isolation-by-distance patterns are present (Frichot et al., 2013). We ran LFMM 2 using the R package LEA (v 3.9.5, Frichot & Francois, 2015) and the lfmm2() command. The number of latent factors (K=3) was chosen based on the deflation of explained variance along the first three principal components. LFMM cannot handle missing locus data, so we imputed data (LEA::impute() command) using K=3 prior to analysis. The second GEA method, redundancy analysis (RDA), is a multivariate approach in terms of both environmental predictor and the genetic response variables (Capblancq & Forester, 2021). Multivariate genetic analysis may account for polygenic architecture in adaptive traits. We used the rda() command in the vegan package (v. 2.6.4, Oksanen et al., 2022).The imputed genotype file that was created for the LFMM analysis was also used in this analysis for consistency across methods. SNPs were scaled prior to analysis (scale=T). We used a full RDA model with climatic variables only, rather than a partial model, because there were strong correlations among genetic principal components, geographic variables, and climatic variables that strongly reduced the signal (Fig S2) (Capblancq & Forester, 2021). The custom command “rdadapt()” from the accompanying R script was then used to calculate q-values using K=3 (Figs S3, S4).

### Identifying signatures of selection across methods

Combining multiple genome scan analyses improves study power to reject neutrality, but false discovery rates within and among tests must be controlled to account for multiple testing and confounding effects (François, Martins, Caye, & Schoville, 2016). The null hypothesis underlying all genome scan tests is that a locus is not under selection, and all four aforementioned methods use chi-squared distributions of putatively neutral alleles to reject this null hypothesis on a per-locus scale. This shared methodological basis makes it possible to compare results among tests, as long as the significance values are properly calibrated. The first calibration step occurs within methods and is based on inherent test statistics used to reject the null hypothesis. In LFMM, pcadapt, and RDA, test statistics were calibrated via the genomic inflation factor. Inflation calibration is automatically implemented in pcadapt and RDA, but it is manually implemented in LFMM using the lfmm2.test() command. BayPass is calibrated with the population covariance matrix created during core model analysis.

Next we curated a study-wide list of candidate outliers by first converting the calibrated p-values to q-values using the p.adjust() command in the base R stats package with the Benjamini-Hochberg equation (Benjamini & Hochberg, 1995), then applying the same false discovery rate control threshold across tests (François et al., 2016). An ideal false discovery rate threshold balances Type I errors (false positives) and Type II errors (false negatives). We applied a fairly lenient study-wide false discovery rate threshold of 0.01 (i.e. < 1% false positives). Finally, we compared overlap among all four tests to determine evidence strength for true positive outliers.

### Annotating outliers

To determine whether outlier SNPs would change the amino acid produced by its containing codon (i.e. synonymous or nonsynonymous substitution), we again used nucleotide BLAST with the contigs containing the candidate SNPs and then translated amino acids within the top gene hit. A recent study found that both synonymous and nonsynonymous mutations can affect the level of mRNA expression of a mutated gene, and that the effect’s magnitude partially predicted the phenotypic fitness outcome (Shen, Song, Li, & Zhang, 2022). In our study, many BLASTn top gene hits included either frame shifts or premature stop codons before the target locus, calling into question the impact of substitution. Therefore, we reported the substitution type in the full outlier list (Table S1), but we did not differentiate among them in our gene function analysis or discussion. Gene functions were obtained from the UniProt database (The UniProt Consortium, 2023). We used PANTHER to summarize GO results and run a GO-term statistical overrepresentation test, using our starting candidate gene list as the background list (Thomas et al., 2022).

## Results

### Population genetic structure and diversity

We found significant negative correlations between each genetic diversity parameter and latitude (Fig 2), meaning diversity values are highest in low-latitude sampling sites and decrease poleward. For example, the correlation between latitude and expected heterozygosity has a high R-value of −0.87 and significant p-value of 7e-07. On the contrary, there are no consistent significant local elevation trends within paired sites, meaning the locally higher sites do not always have lower diversity. In the north, diversity values tend to be greater in high-elevation sites (e.g. sites 2 & 4), but in the south they tend to be lesser in high-elevation sites (sites 15, 17, 19). The same patterns hold true for nucleotide diversity, which had overall the weakest correlation with latitude (R = − 0.57, p = 0.0082).

**Figure 2.**
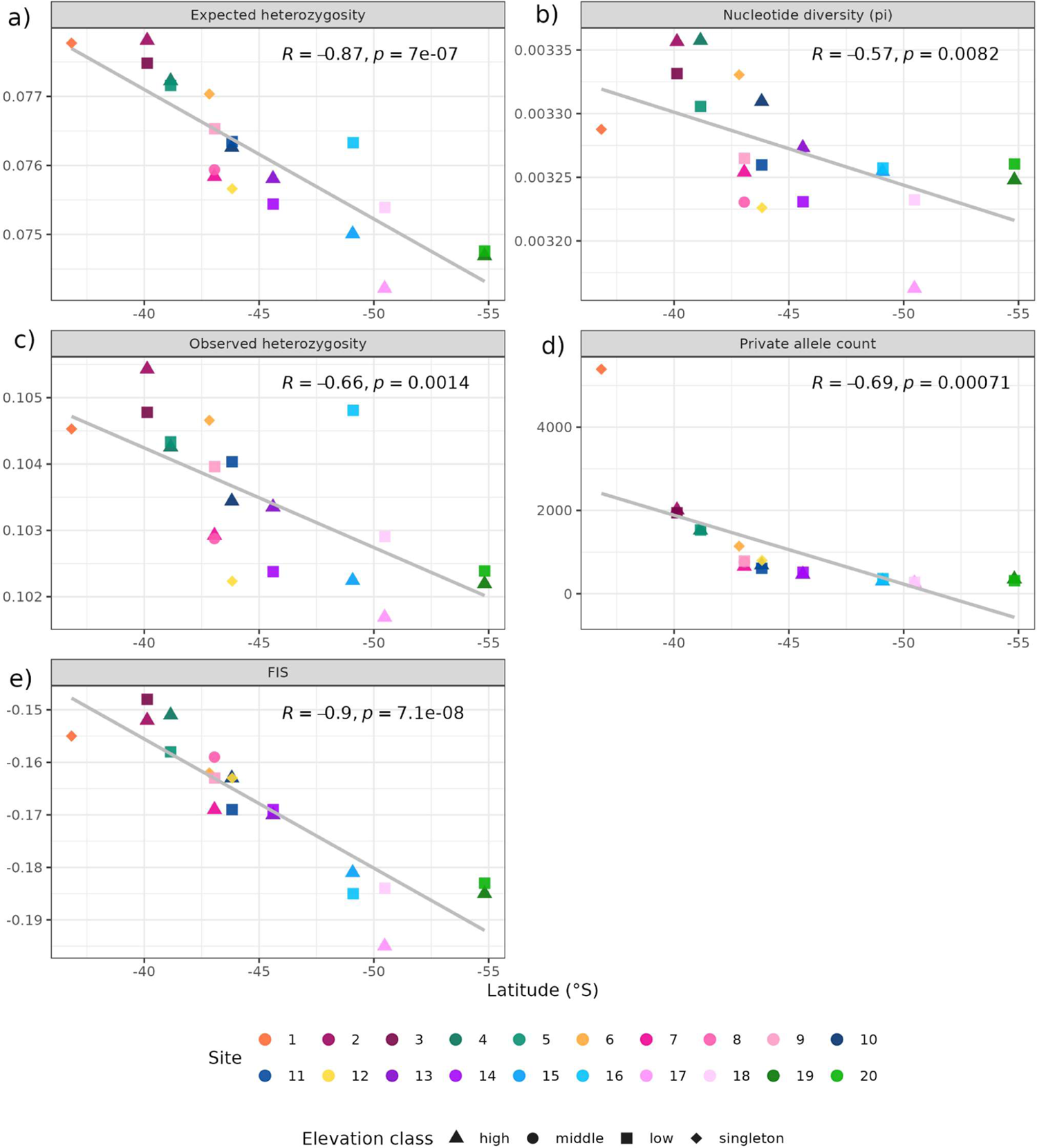
Genetic diversity statistics and their correlations with latitude. a) expected heterozygosity, b) nucleotide diversity including all invariant called sites, c) observed heterozygosity, d) rarefied private allele count, and e) fixation index F_IS_. Colors indicate site number and shapes indicate relative elevation class (high, middle, and low within paired localities, or singleton). R-squared and p-values for linear regression models are included in each graph.

Observed heterozygosity per sampling site was always greater than expected and therefore there is heterozygote excess (negative *F*_IS_), as can be expected from a self-incompatible outcrossing species. Pairwise subpopulation differentiation (*F*_ST_) values were all less than 0.036 (Fig 3) and had a mean of 0.0102. Paired sites generally had the lowest pairwise *F*_ST_ values, with most ranging from 0.00024 (sites 7 vs. 8) to 0.0025 (19 vs. 20). The notable exception is sites 17 and 18, which have a value of 0.0072. Singleton locality values in relation to all sampling sites ranged from 0.0025 to 0.0359. Site 1 (Epulaufquen) consistently had the highest pairwise values with all other populations (range 0.0196 - 0.0359). Epulaufquen also had the greatest endemic diversity (number of private alleles), and then the values sharply decreased poleward.

**Figure 3.**
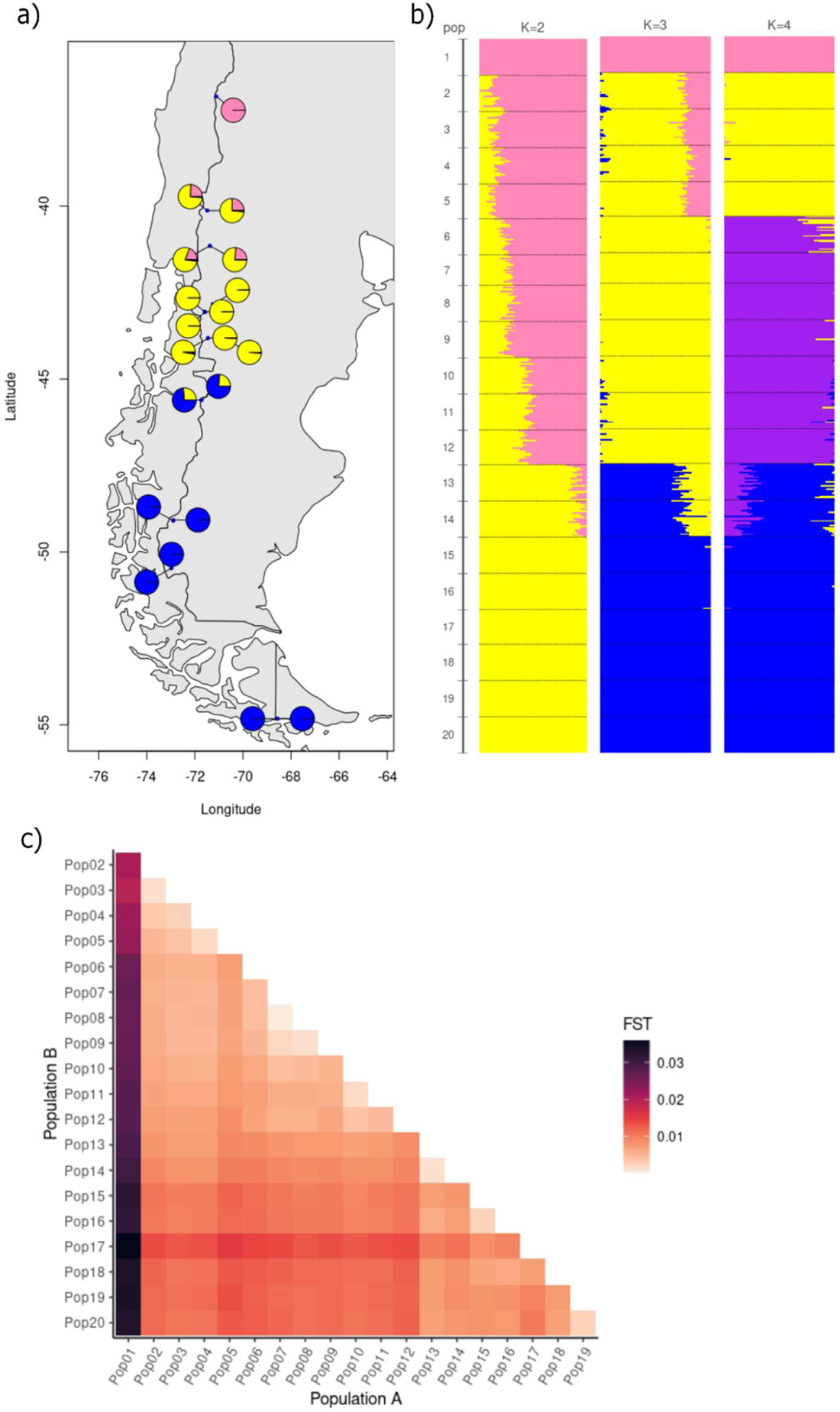
Population genetic structure of *Nothofagus pumilio*. a) Average cluster per sampling site using K = 3 results from ADMIXTURE. b) Individual ADMIXTURE plots for K values of 2, 3, and 4, ordered top to bottom from north (sampling site 1) at the top to south (20) at the bottom. Each line indicates one individual and colors indicate population clusters. (c) Pairwise genetic difference between and among sampling sites as assessed with Weir and Cockerham pairwise F_ST_ values. Sampling sites are ordered from North (site 1) at top and left to South (20) at bottom and right. Color indicates F_ST_ value, from small (light orange) to large (dark purple).

Population structure is also oriented along the latitudinal gradient, although the exact number of genetic clusters is ambiguous. According to cross-validation values in ADMIXTURE, the optimal number of genetic clusters (K) is 2. However, principal component and snmf (LEA package) analyses suggested that the optimal number is 3 (Fig 4a). We present K values from 2-4 (Fig 4b), since all are informative about the hierarchical population structure (Meirmans, 2015). Across K-values, a break consistently occurs between sites 12 and 13 (i.e. between 43.8 - 45.6 °S), with admixture appearing in sites 13 and 14. At K=3, sites 2-5 show admixture, and at K=4 this region becomes its own cluster, with a break between sites 5 and 6 (41.2 - 42.8 °S). The singleton northernmost site 1 (Epulaufquen) also isolates into its own cluster at K=4.

**Figure 4.**
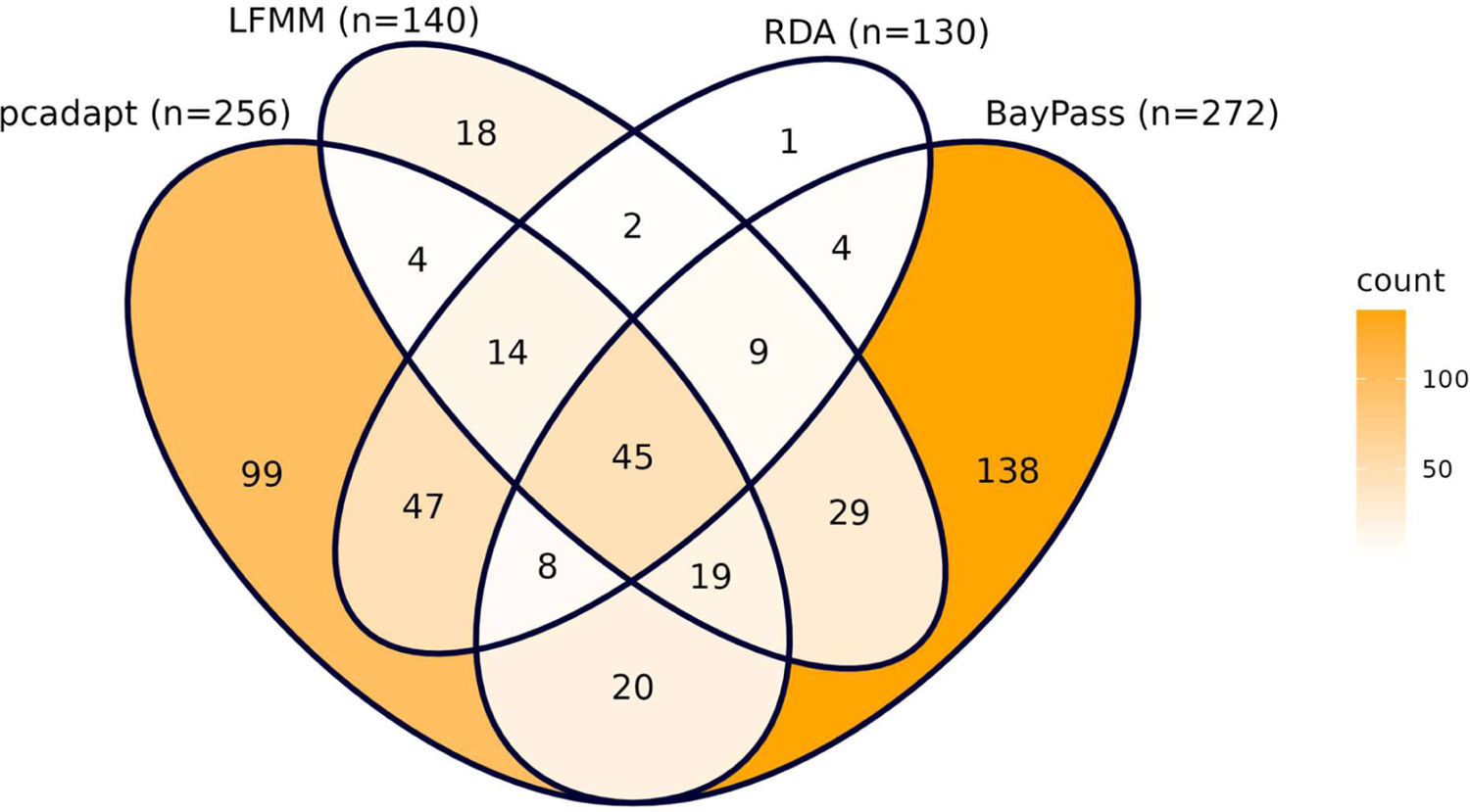
Overlap among significant outlier SNPs found by each genome scan method. Numbers inside each Venn section indicate the number of SNP(s) found by the method(s), and color also indicates count, from low (white) to high (dark orange). Total number of SNPs found by a method is shown in parentheses after the method name.

### Climate conditions per site

Climate covariates also show strong gradients along the latitude axis, and many have strong within-locality gradients (Fig S1). Higher-elevation sites experience more frost days and greater total annual precipitation, although the majority is received as snow in winter (Veblen et al., 1996, Fig S1b & 1d). Northern localities generally have higher temperatures and annual precipitation amounts than central and southern localities, although the low-latitude Epulaufquen locality (site 1) is among the drier localities. The El Chaltén (sites 15-16) and El Calafate (17-18) localities are directly adjacent to the Southern Patagonian icefield, where most precipitation falls as snow even in summer, and therefore they are among the coldest and driest sites overall. The southernmost locality Ushuaia (19-20) has a maritime climate, with relatively warmer temperatures and higher precipitation than other poleward localities. The two geographically marginal singleton localities in the central portion of the range (6-12) have lower precipitation and more frost days than their counterpart paired localities at similar latitudes (sites 7-9 and 10-11, respectively), demonstrating that they are also environmentally marginal. A PCA indicated 81% of the variance can be described by two axes, the first mainly comprising annual precipitation and day length in January, and the second comprising number of frost days and growing season precipitation (Fig S1f).

### Genes containing SNP outliers

A total of 457 SNPs in 329 contigs (5.2% of analyzed SNPs) were identified as outliers by at least one genome scan method under a false discovery rate threshold of 1% (Table S1). BayPass identified the greatest number of outliers (n=272) and also had the greatest number of unique outliers (Fig 4). RDA identified the fewest outliers (n=130), and only one outlier was unique to this analysis. Of the RDA outliers, 114 were also found by the other multivariate-genetic method, pcadapt. Meanwhile, pcadapt found 99 unique outliers that were not identified by any other method.

Of all outliers, 201 SNPs were identified by at least two methods, and 45 of these were identified by all four algorithms (Fig 4). These two outlier lists will be called moderate-evidence and strong-evidence outliers (i.e. for being true positives), respectively. The majority of moderate- and strong-evidence outliers are located within coding regions of annotated target genes (125 annotated of 201 total outliers, Table S1). The GO enrichment analysis with PANTHER indicated that no gene functions were significantly overrepresented in relation to the background starting gene list.

BayPass is a univariate method for both predictor and response variables, so it is possible to identify individual predictors per SNP (Fig 5). As an aside, LFMM could have been used in univariate-environment mode, but we chose to use its multivariate configuration to reflect the real-world dimensionality of environmental covariates. Day length in January and isothermality were significantly associated with the most outliers (n=113 and 106, respectively). Among the 45 strong-evidence outliers, all but 3 were associated with one or both of these covariates. Growing season precipitation was significantly associated with the fewest outliers (n=24). Thirty-eight of the BayPass outliers were associated with more than one covariate, and one SNP was significantly associated with all five covariates. An example of a strong-evidence SNP that associated with more than one environmental covariate is sequence “chain_2392, locus 1061” (Table 2). The allele cline is plotted against the three environment clines with which it associates in Figure 6. The allele frequency cline also demonstrates within-locality allele differences, most noticeably in El Calafate (sites 17-18). This SNP is located within the MYC2 gene, a transcription factor that may be involved in light signaling pathways and abscisic acid signaling pathways, which is related to drought stress response (Yadav, Mallappa, Gangappa, Bhatia, & Chattopadhyay, 2005).

**Figure 5.**
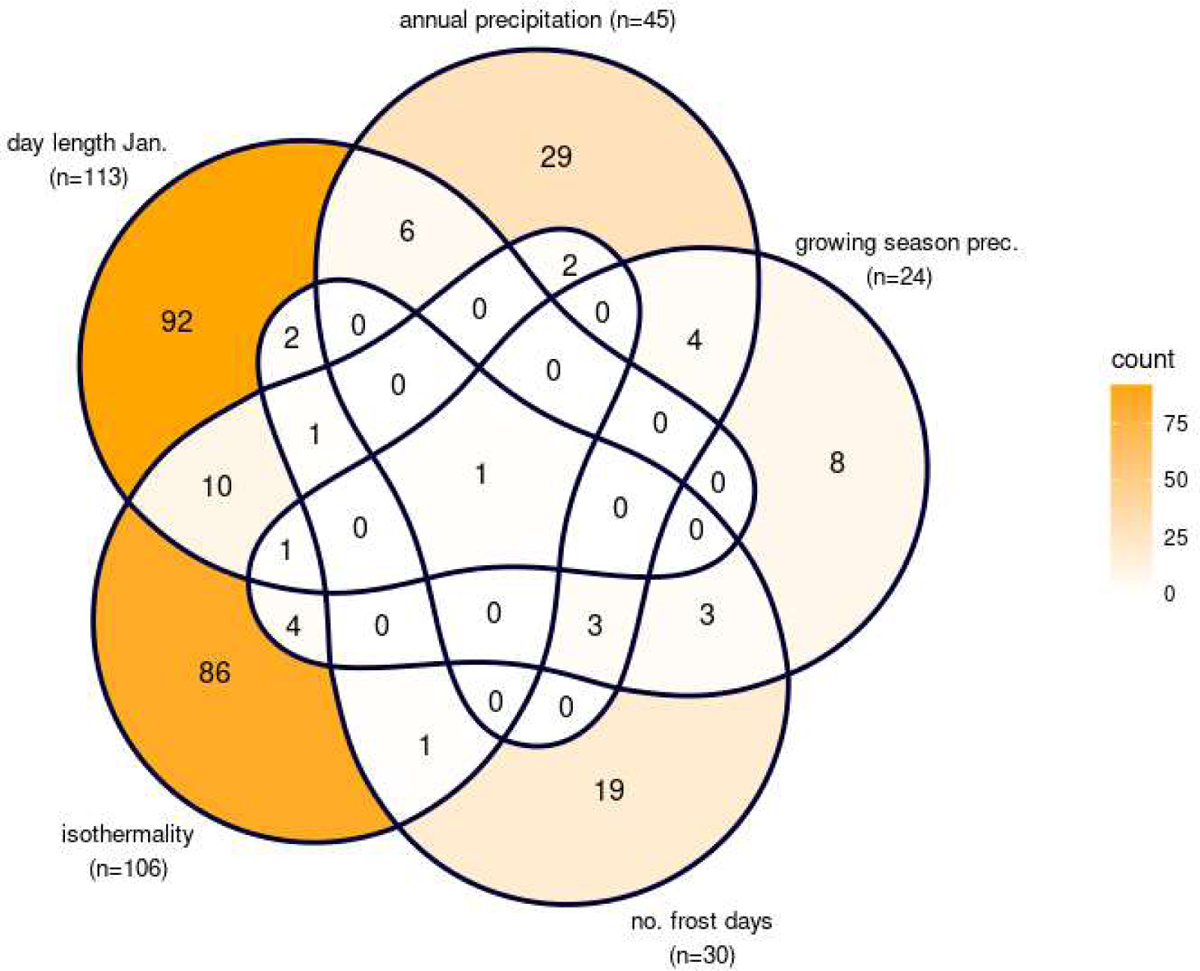
Number of outlier SNPs significantly associated with individual covariates in the univariate method BayPass. Threshold for significance is Bayes Factor > 10. Color indicates count, from low (white) to high (dark orange). Total number of associated SNPs is shown in parentheses after the covariate name. Numbers inside each Venn section indicate the number of SNP(s) associated with the covariate(s).

**Figure 6.**
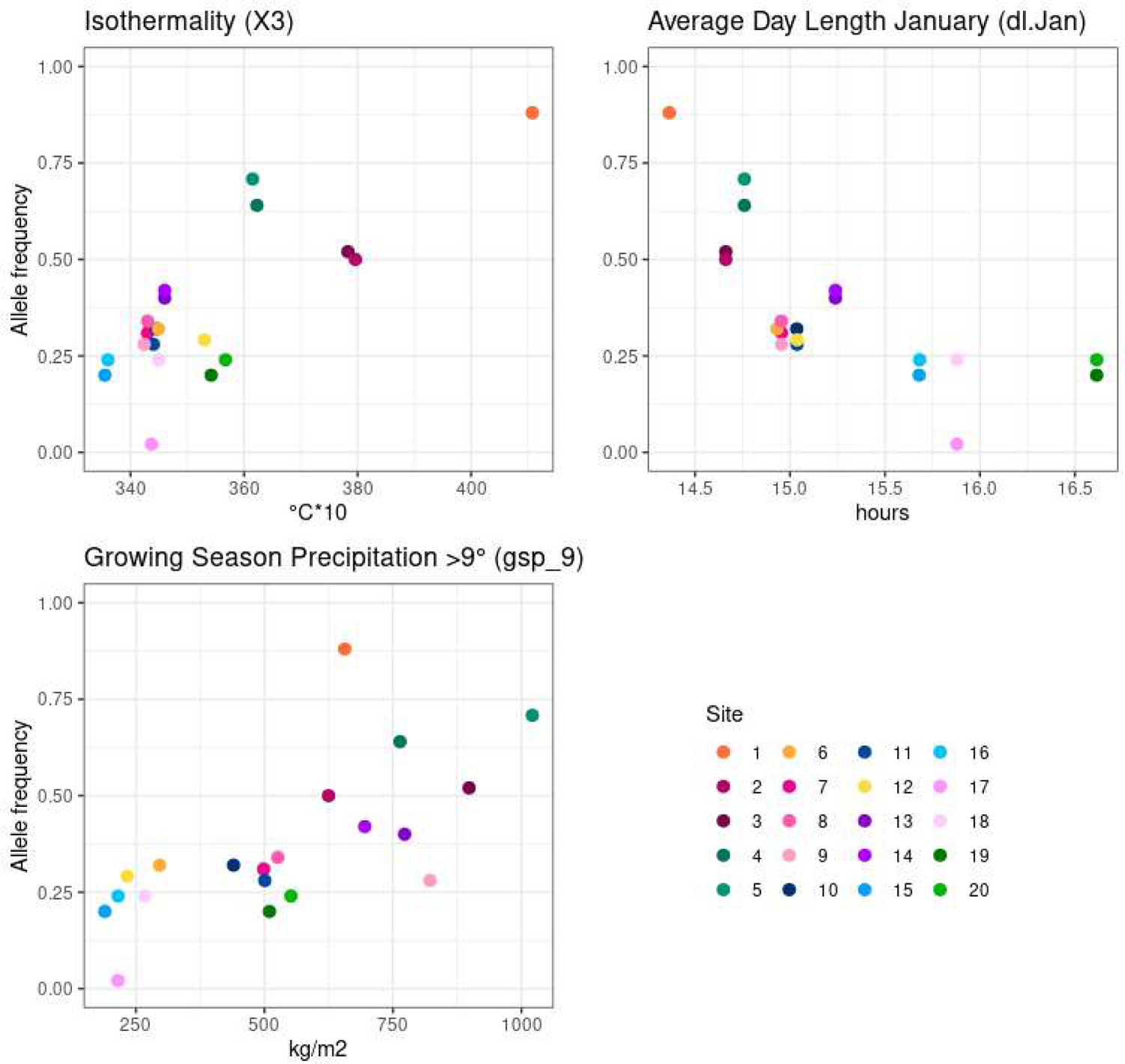
Example allele frequency clines for a single SNP locus that is significantly associated with multiple covariates. The sequence is “contig “chain_2392,” nucleotide position 1061. The covariates are isothermality, average day length in January, and growing season precipitation. Colors indicate sampling site. This SNP is located within gene MYC2. Allele frequencies are not corrected for population genetic structure.

**Table 2.**
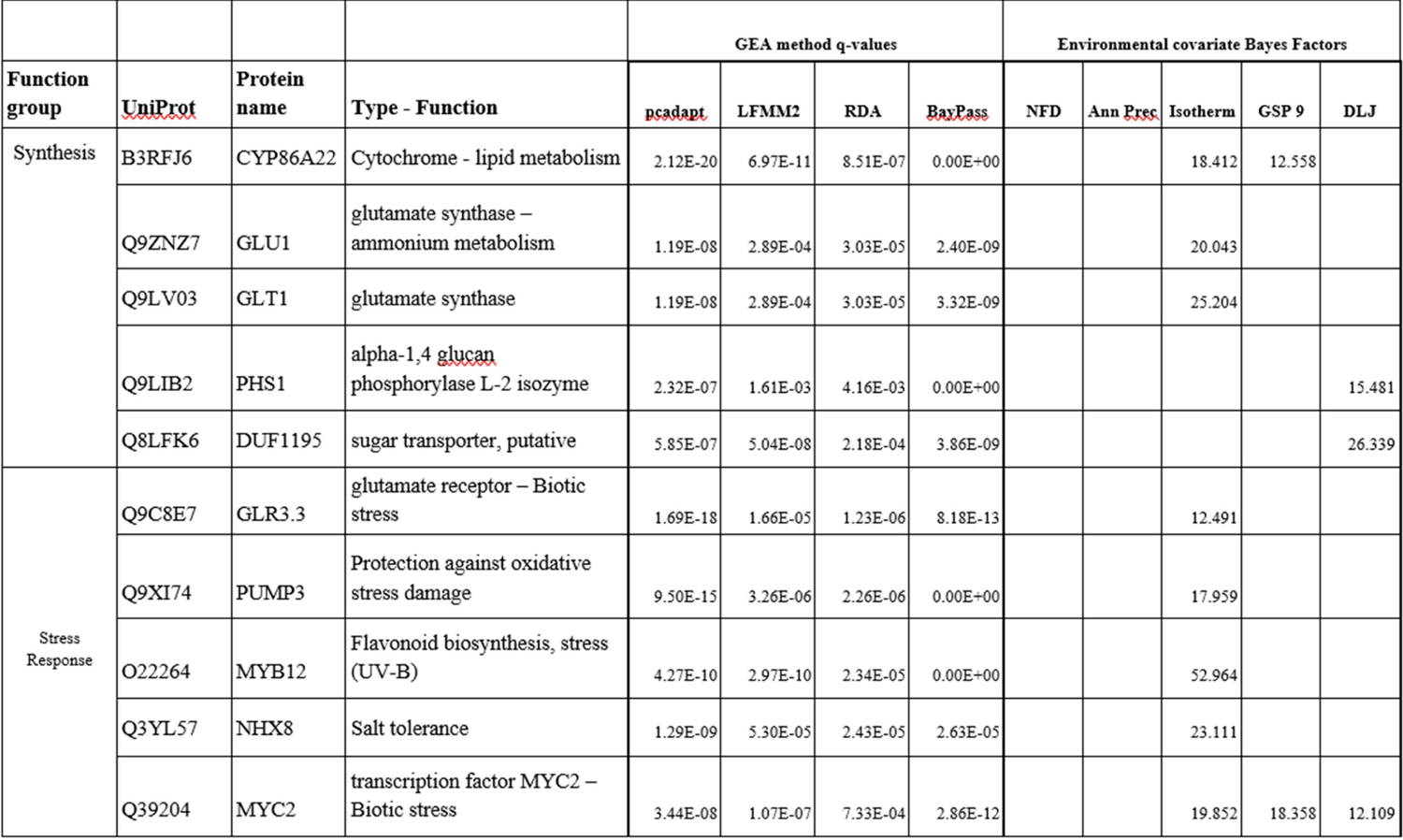
Gene function summary for a subset of the strongest evidence outliers within the synthesis-metabolism and stress response protein function groups. UniProt accession code, prottein name, and function were sourced from UniProt database. Q-values for each GEA method are shown, as are all significant Bayes Factor values for environmental covariates that had significant q-values in BayPass. For further details on these and other outliers, see Supplemental Table 1.

## Discussion

Climate change will likely disrupt relationships among environmental covariates, thereby exerting unique selection pressures on trees such as the Andean foundation species *Nothofagus pumilio*. We used a landscape genomics approach to determine which genes show signatures of adaptation and which environmental factors might be influencing selection. In particular, we investigated environmental covariates related to temperature, day length, and precipitation, both as univariate and multivariate factors, since interplay among covariates can affect biological processes such as phenology and drought response. We found that population structure and genetic diversity in *N. pumilio* are mainly structured along the major north-south spine of the Andes, likely due to past glacial cycles. Temperature and day length variables were significantly associated with the greatest numbers of outlier SNPs, while precipitation variables were associated with fewer. However, many outliers were identified only by multivariate analyses or were associated with more than one variable, suggesting genes are responding to a combination of environmental covariates. Climate change may have a dire impact on *N. pumilio* survival and adaptation, particularly if it decouples relationships among environmental selection pressures to which these genes are currently adapted.

### Latitude-oriented population structure and genetic diversity patterns reflect glacial oscillations

The latitudinal orientation of the hierarchical population structure (Fig 2) is likely due to glacial oscillations. Our population structure analyses (K = 2–4, Fig 4) indicate the first major cluster division occurs at mid-latitudes between 43–45 °S (sites 12 and 13). Previous neutral marker studies also found evidence for two geographically segregated chloroplast and microsatellite lineages, namely a northern and a southern clade (Mathiasen & Premoli, 2010; Mattera, Pastorino, Lantschner, Marchelli, & Soliani, 2020; Soliani, Gallo, & Marchelli, 2012; Soliani et al., 2015). Those analyses identified the major division slightly further north, near 42 °S (Mathiasen & Premoli, 2010; Mattera et al., 2020), or between 42 - 44 °S (Soliani et al., 2015). At K = 4, we also observed a division there, specifically between 41.1 - 42.8 °S (sites 5 and 6). This phylogenetic divide has also been observed in other regional taxa (e.g. Sersic et al., 2011) and has been attributed to divergent glacial patterns. Glaciers north of 41 °S were alpine-style and restricted to valleys, while glaciers south of 45 °S formed a more continuous ice sheet (Glasser, Jansson, Harrison, & Kleman, 2008). Palynological and genetic data indicate *Nothofagus* species responded to these glaciations by migrating towards more favorable northern latitudes (Villagran, 1990) and by retreating to refugia in various parts of the range (Markgraf, 1993).

The distribution of refugia and subsequent postglacial expansion have helped shape the genetic landscape of *N. pumilio*. In general, high heterozygosity is expected in regions near refugia (Petit et al., 2003; Roberts & Hamann, 2015) and also in admixture zones along recolonization routes where secondary contact occurred (Soliani et al., 2015). We found higher heterozygosity values in the north, providing evidence for more northern refugia than southern. However, we found no elevated heterozygosity near the mid-latitude population cluster divisions, even though admixture in sites 13 and 14 (El Triana), which suggests this region was a secondary contact zone. It is possible that this lack of elevated heterozygosity is due to unobserved evolutionary forces such as genetic drift or bottlenecks. Assuming a similar mutation rate across populations, higher nucleotide diversity in the north may also reflect larger historical effective population sizes there (Nei & Takahata, 1993). This would be a likely consequence of northward migration patterns and denser refugia distribution. Greater endemic diversity (private alleles) in the north also support the idea of enduring populations that may have been temporarily isolated from other populations. Notably, southern sampling sites also had at least 250 private alleles each, supporting the claim that there were multiple southern refugia in previous Glacial Maxima (Paula & Leonardo, 2006; Andrea C Premoli, Mathiasen, & Kitzberger, 2010). Further evidence regarding postglacial expansion patterns comes from gene flow and migration speed estimates. Species-specific gene flow information is limited, although it has been postulated that wind-dispersed pollen can travel 10-100 times further than the anemochorous seeds (Mathiasen & Premoli, 2010). High male-gametic gene flow could explain our low overall population differentiation values, which have a mean of 0.0102 (Fig 3). On the other hand, a prior estimate of migration speed suggests that the maximum postglacial expansion distance following the Last Glacial Maximum is no more than 800 km (Mathiasen & Premoli, 2010).

There were no significant range-wide elevation patterns among diversity statistics, although local gradients and regions showed trends. High-elevation sites in poleward localities generally had lower diversity, which aligns with previous studies that found high-elevation populations of *N. pumilio* have reduced polymorphism due to recent postglacial expansion, genetic drift, and/or inbreeding (A C Premoli, 2003). Suboptimal climate conditions at higher elevations may also be at play (Mathiasen & Premoli, 2013), for example the increased number of frost days (Fig S1e). In contrast, high-elevation sites in low-latitude localities generally had greater diversity. A possible explanation is that our “high-elevation” sites were located below local treelines, meaning they could be more accurately classified as intermediate elevation. Meta-analyses have shown intermediate elevation areas have high genetic diversity due to locally optimal environmental conditions and larger effective population sizes (Ohsawa & Ide, 2008).

A peculiar case is observed in the northernmost site, Epulaufquen (site 1), which had the highest number of private alleles and ubiquitously high pairwise *F*_ST_ values, but also had relatively low observed heterozygosity and nucleotide diversity (Fig 2). Despite strong genetic differentiation from other sites (Fig 3c), this site only segregates into its own population cluster at K=4 (Fig 3b). This may be due to its relatively small sampling size (e.g. Rosenberg et al., 2002) in comparison to the paired localities, or there is relatively stronger differentiation between north and south clusters at the range-wide level. Epulaufquen is an ecologically and topographically unique site, and related species with populations there such as *Nothofagus obliqua* also display distinct genetic and morphological characteristics (Azpilicueta et al., 2014). The forest is located on the valley floor and is surrounded by mountain peak barriers that likely impede migration and gene flow, which may explain the high *F*_ST_ values and private allele count. Epulaufquen is near the species’ current low-latitude range margin, which are usually areas that experience higher temperatures and less precipitation in comparison to the rest of the range and are thus exposed to increased drought risk (Hampe & Petit, 2005). This locality may harbor alleles that are adapted to extreme conditions, but it is also possible that this locality will not remain a suitable habitat for *N. pumilio* under warmer conditions.

### Evidence sources

Univariate and multivariate genome scan methods provide different but equally valuable information about signatures of local adaptation. Univariate genetic methods can identify large-effect alleles that may be under strong selection, and univariate environment methods can identify the drivers exerting the strongest selection pressures. Meanwhile, multivariate genetic methods can identify small-effect alleles, and multivariate environment methods can characterize interplay among environmental factors. While multivariate environment methods may be more realistic, an important caveat is that they combine covariates into principal components, which are inherently difficult to interpret (Rellstab et al., 2015). In this case, it can be more informative to examine the biological functions of the outlier-containing genes. Therefore, we used a combination of univariate and multivariate results and the biological processes of outlier-containing genes to characterize selection pressures.

PCAdapt identified 99 outliers that were not indicated by any of the GEA analyses. Population differentiation methods may have more power than GEA analyses when demographic history has caused collinearity between neutral allele frequencies and environmental clines (Lotterhos & Whitlock, 2015), for example in a post-glacial orographic habitat like Patagonia. This explanation is supported by our results, which indicate that genetic diversity statistics (Fig 2), population structure (Fig 3), and environmental covariates (Fig S1) all had strong relationships with the latitude cline. However, allele frequencies do differ among sites within localities (Fig 8), demonstrating that latitude is not the only axis along which adaptation occurs. Therefore, it is possible that these 99 putatively adaptive regions are associated with unobserved climatic factors or they may be influenced by factors beyond climate (e.g. Meirmans, 2015). The pcadapt-unique outliers may be worth investigating in further studies.

### Temperature and day length most important environmental covariates

In the univariate GEA test BayPass, we found notably more SNP outliers that significantly associated with a temperature (n=133) and/or photoperiod (n=113) covariate than with a precipitation covariate (n = 61) (Fig 6, Table S1). Temperature and photoperiod parameters may have a stronger effect than precipitation does, an interpretation that aligns with some previous studies of *N. pumilio* adults and seedlings. In adults, significant correlations were found between radial growth and temperature but not precipitation (Castellano, Srur, & Bianchi, 2019). In seedlings, temperature was also a stronger factor than air humidity for mortality rate along elevation clines (Cagnacci et al., 2020). Our study is the first to explicitly assess photoperiod in relation to *N. pumilio* genetic adaptation and the 113 day length-associated outliers suggest that photoperiod has a strong effect. Since empirical information is limited for *N. pumilio*, we turn to supporting evidence from other tree species. Delayed spring bud-burst in response to short photoperiod has been observed in related late-successional species *Fagus sylvatica* and *Quercus petraea* under common garden conditions (Basler & Körner, 2012; Vitasse & Basler, 2013). Similarly, autumnal dormancy in perennial plants is largely initiated by the environmental cues of shortened photoperiod and low temperature (Howe et al., 1995; Singh et al., 2017); for example in *Populus* species, dormancy-related genes (phytochromes) have shown stark latitude clines (Ingvarsson, Garc\’\ia, Hall, Luquez, & Jansson, 2006). These results highlight the importance of temperature, photoperiod, and their interaction in regards to phenology and adaptation.

Fifteen outliers are associated with both day length and at least one temperature covariate, and the containing genes are predominantly related to stress response (Table S1). This temperature-photoperiod overlap value is somewhat lower than expected, considering that both temperature and photoperiod clines are mainly oriented along the latitude axis (Fig S1), but covariate pruning is an important consideration when examining these results. We chose isothermality and number of frost days for the temperature variables due to their biological relevance, but also in part because their correlations with January day length were moderate (R = −0.44 and R= −0.39, respectively; Fig S1). However, some pruned temperature parameters had much stronger correlations with day length (R > 0.9, data not shown). Environmental associations must always be carefully interpreted (e.g. Rellstab et al., 2015), and outliers that we found to be associated with photoperiod might actually be associated with unobserved but highly correlated temperature variable(s) instead of, or in addition to, photoperiod. In any case, unobserved relationships may well be disrupted by global climate change, meaning the outlier-containing genes will still experience different selection pressures than those under which they evolved.

A smaller number of precipitation-associated outliers were also identified (n=61), and the containing annotated genes have diverse functions including metabolism, growth, and stress response (Table S1). However, only one of these annotated genes is directly related to drought response (probable protein phosphatase 2C 24). A notable point is that 11 annotated outlier-containing genes were assigned Gene Ontology terms related to water deprivation response, but the SNPs were not significantly associated with either precipitation covariate (e.g. genes NCED1, BFRUCT3). Possible reasons for the lower count of precipitation-associated outliers may include sampling locations or *N. pumilio* biology. Regarding the study design, our sampling area encompassed the drier portion of the species’ range, namely in Argentina. Our sampling sites received between 380-1600 mm mean total annual precipitation between 1979-2013, but the species also inhabits Chilean locations that received up to 5500 mm of mean precipitation during those years (Figs 1, S1). It is possible that our sampled range was insufficient to capture more precipitation-related signatures of adaptation. From a biology standpoint, common garden studies performed on young *N. pumilio* that were collected along local precipitation or elevation gradients consistently show trait differentiation in water use and morphology when those plants are grown under drought conditions, but there is little consensus about whether genetics or phenotypic plasticity is responsible. Some studies suggest a genetic basis (Ignazi, Bucci, & Premoli, 2020; Soliani & Aparicio, 2020), others suggested that responses are plastic (Ivancich et al., 2012), and still others found supporting evidence for both mechanisms (Mathiasen & Premoli, 2016; Andrea C Premoli & Brewer, 2007; Soliani et al., 2012). Phenotypic plasticity is advantageous when physical conditions are highly variable, for instance in northern Patagonian locations with a Mediterranean climate (Villalba et al., 2003). Plasticity can allow plants to evade temporary suboptimal conditions, but sidestepping the selection pressures required for adaptation means fewer loci may show adaptive signatures. Joining the common garden evidence with our genome scan results, it seems likely that water use and drought response may be complicated processes that implicates many biological processes and could also incorporate plasticity.

The critical question of how interplay among environmental covariates affects selection has complicated answers. Interactions between precipitation and temperature have been noted as important for *N. pumilio* survival and radial growth (Lara et al., 2001) as well as seedling establishment patterns at alpine treelines (Daniels & Veblen, 2004). Interactions among day length and temperature are also known to be important in the annual growth cycle and phenology (Singh et al., 2017). Of the 272 BayPass outliers, 38 were associated with more than one univariate environmental factor. We explicitly assessed environmental interplay through two multivariate environmental analysis methods, which identified 40 annotated SNPs that were not identified by the univariate method. While synthetic variables used in multivariate analyses can represent environment space more realistically, they are inherently difficult to interpret biologically. Therefore, it is informative to examine the biological functions of the outlier-containing genes.

### Biological functions of outliers mainly related to stress and metabolism

The tradeoff between growth and survival is a fundamental challenge for plants, particularly when they undergo stress events such as high temperature or drought. Nearly half of all annotated outliers were located in genes related to stress response (n=38, 21%) or metabolism-catabolism synthesis (n=43, 24%) (Table S1). A previous study regarding *N. pumilio* transcriptome expression under heat stress found that genes related to photosynthesis and carbon metabolism were down-regulated under heat treatment, while stress response genes were up-regulated (Estravis-Barcala et al., 2021). Many of our outlier-containing genes were similar in form or function to those identified in the heat stress study. For example, we found 6 related outlier-containing genes related to the ABA signaling pathway, which was up-regulated by heat stress. ABA is produced under water deficit and confers tolerance to water and salt stress (Abe et al., 2003). Among our ABA-related outliers were CBL9 and CCD1, which had no univariate associations, and MYC2, which is also a regulator of light and jasmonic acid (Yadav et al., 2005) and was associated with all three major variables (Fig 8). Taken together, this provides evidence that the ABA pathway experiences diverse selection pressures. Another family of genes, the WRKY transcription factors, contained multiple outliers and was also up-regulated under heat stress (Estravis-Barcala et al., 2021). While the exact genes differed between studies, the genes’ functions were comparable, namely defense response to drought or pathogens (WRKY 4, associated with day length and temperature, and 33, associated with day length).

Nested within the growth-survival tradeoff is phenology, which also has implications for reproduction and future adaptation. Growth and development comprised 16 genes (9%) and included phenological functions such as flowering time. PHYA is a critical component for flowering time expression through light perception (Yanovsky & Kay, 2002) and a PHYA homolog in *Populus* trees has also been linked to autumnal growth cessation and bud set (Böhlenius et al., 2006). Our results indicate this gene is under selection in *N. pumilio* and is associated with day length. Other flowering time genes, like PFT1, are not associated with any covariates. In the closely-related species *Fagus sylvatica*, spring bud burst phenology is also photoperiodically controlled (Vitasse & Basler, 2013). A disconnect between day length and temperature could mean a longer growing season that trees could not take advantage of, particularly those at higher elevations whose growing-season window is already condensed (Barrera, Frangi, Richter, Perdomo, & Pinedo, 2000).

### Synthesis and outlook

Climate change could decouple photoperiod from contemporary temperature and precipitation patterns, thus placing novel selection pressures upon genes related to phenology and stress response. Climate change has already exacerbated ENSO effects, including acute drought stress, and the Patagonian precipitation gradient is expected to sharpen further (Barros et al., 2015). Populations of *N. pumilio* are adapted to current specific patterns and combinations of environmental variables, and as these patterns and combinations shift, the adaptations could become maladaptations. A promising avenue for predicting climate change impact is to quantify genomic offset (reviewed in Capblancq, Fitzpatrick, Bay, Exposito-Alonso, & Keller, 2020), which characterizes mismatch between extant allele compositions and those that would be required under future conditions. In order to evaluate genomic offset under no-analog conditions, it must first be established that local adaptation clines are associated with environmental clines, as we have shown here.

We found evidence that patterns of local adaptation in *N. pumilio* are associated with temperature, day length, and, to a lesser univariate extent, precipitation. Individual genes related to stress response and development were often associated with temperature and day length. This supports our prediction (i) regarding growth and stress response genes being correlated with temperature and photoperiod. However, against our prediction (ii), genes related to drought response were only rarely associated with precipitation covariates, and were more likely to be associated with temperature and/or day length. This suggests a more complex response to drought than precipitation and genetic factors alone. These results have many implications under future climate change. Cataloging extant genetic diversity may help us identify ideal candidate genes and populations for assisted gene flow or migration, and identifying the most influential local selection drivers may help predict response to climate change.

## Conclusion

Revealing genetic adaptations to climate is a critical exercise for forests in the face of climate change, especially if relationships among environmental variables become decoupled. Here, we investigated relationships among environmental factors and allele frequencies of outlier SNPs in candidate genes in the Andean foundation species *Nothofagus pumilio*. We found that population structure and overall genetic diversity have strong relationships with latitude. Temperature and photoperiod covariates were associated with the greatest number of outlier-containing genes, while precipitation was associated with fewer, even among genes related to drought response. Our results suggest that stress response and catabolism-metabolism may be under tighter genetic control than drought response traits, which could be more plastic. This has great relevance for this species’ ability to survive and thrive into the future.

## Supplemental materials

Supplemental Document 1

