## Supplemental Document 1 for "Temperature and day length drive local adaptation in the Patagonian foundation tree species *Nothofagus pumilio*"

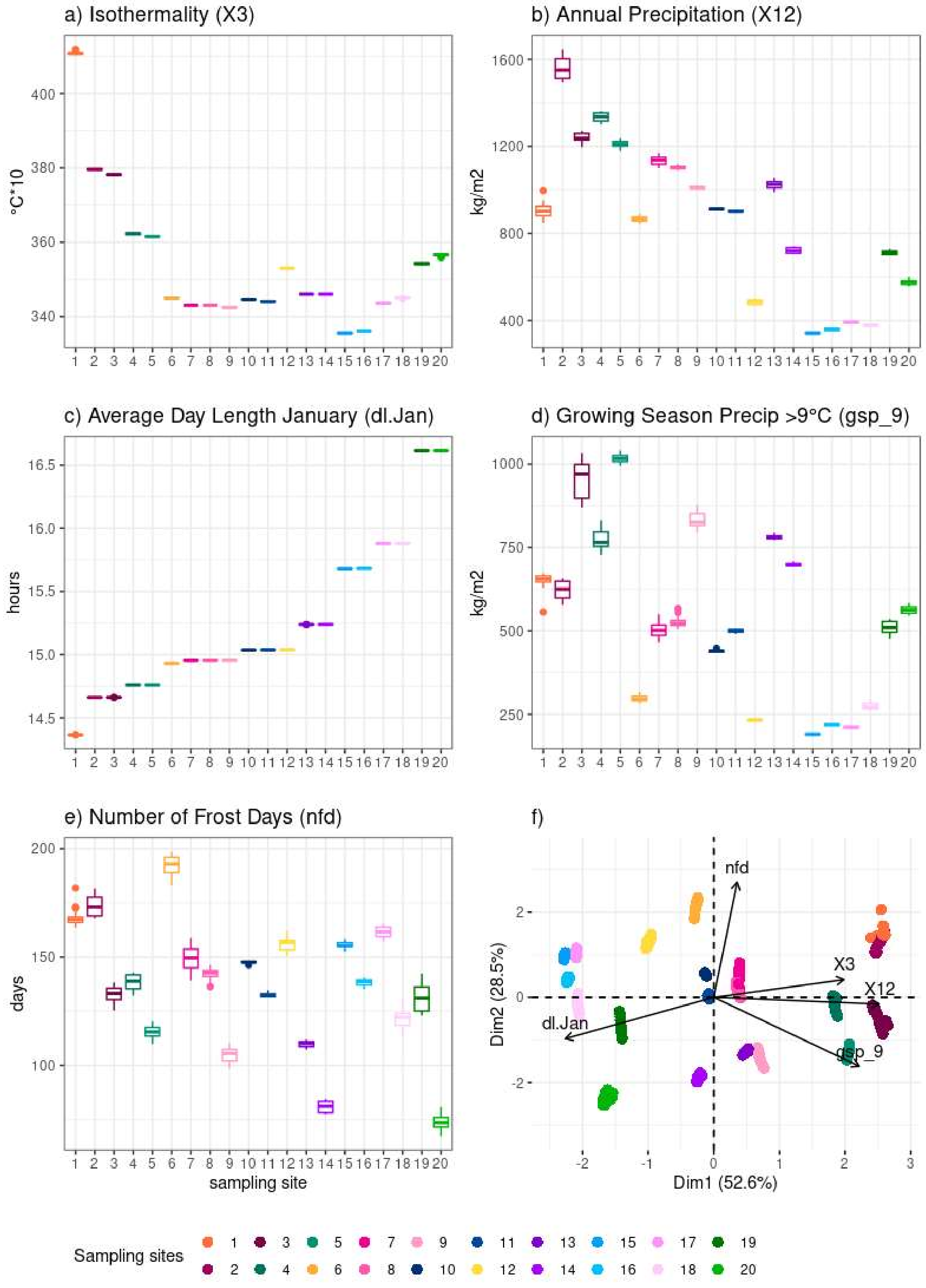


**Figure S1. Boxplots of CHELSA climate conditions per site**. X axis shows the sampling site, also shown in color for comparison with plot f). a) isothermality, b) annual precipitation, c)average day length in January, d) growing season precipitation when temperature is greater than 9°C, e) number of frost days. Finally, f) is a PCA of all five climate conditions at individual tree coordinates.

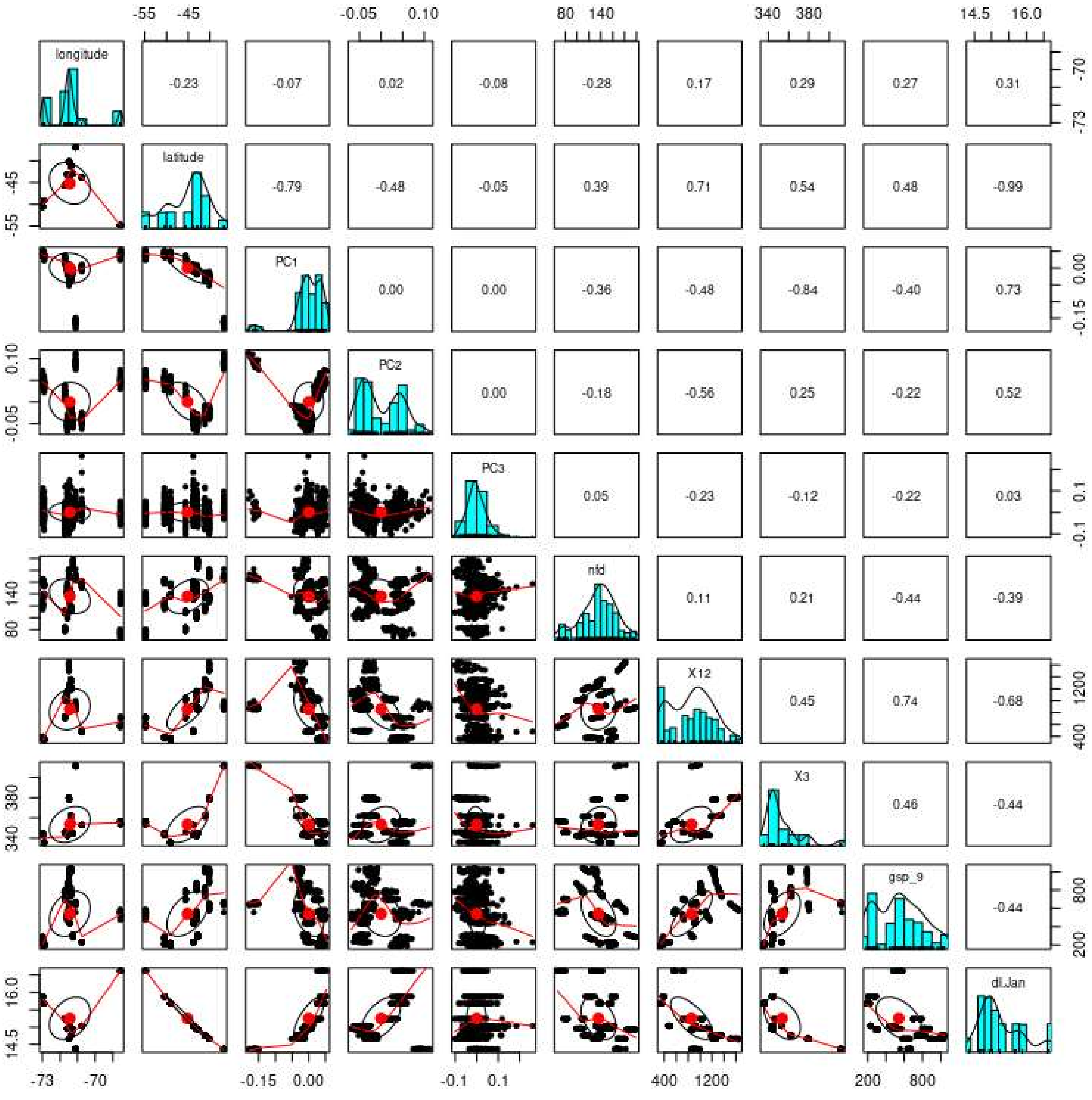


**Figure S2. Correlations among geography, the first three genetic principal components, and the five chosen environmental covariates after pruning highly-correlated environmental covariates.** Graphical representations are shown below the diagonal, numeric values are shown above. Histograms along the diagonal show value distribution within that parameter.

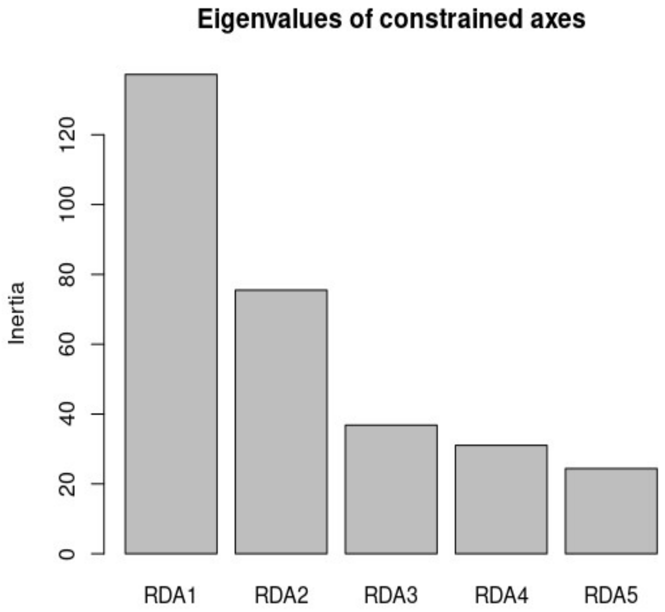


**Figure S3. Eigenvalues of constrained axes in the RDA analysis.** We chose K=3 as the number of axes.

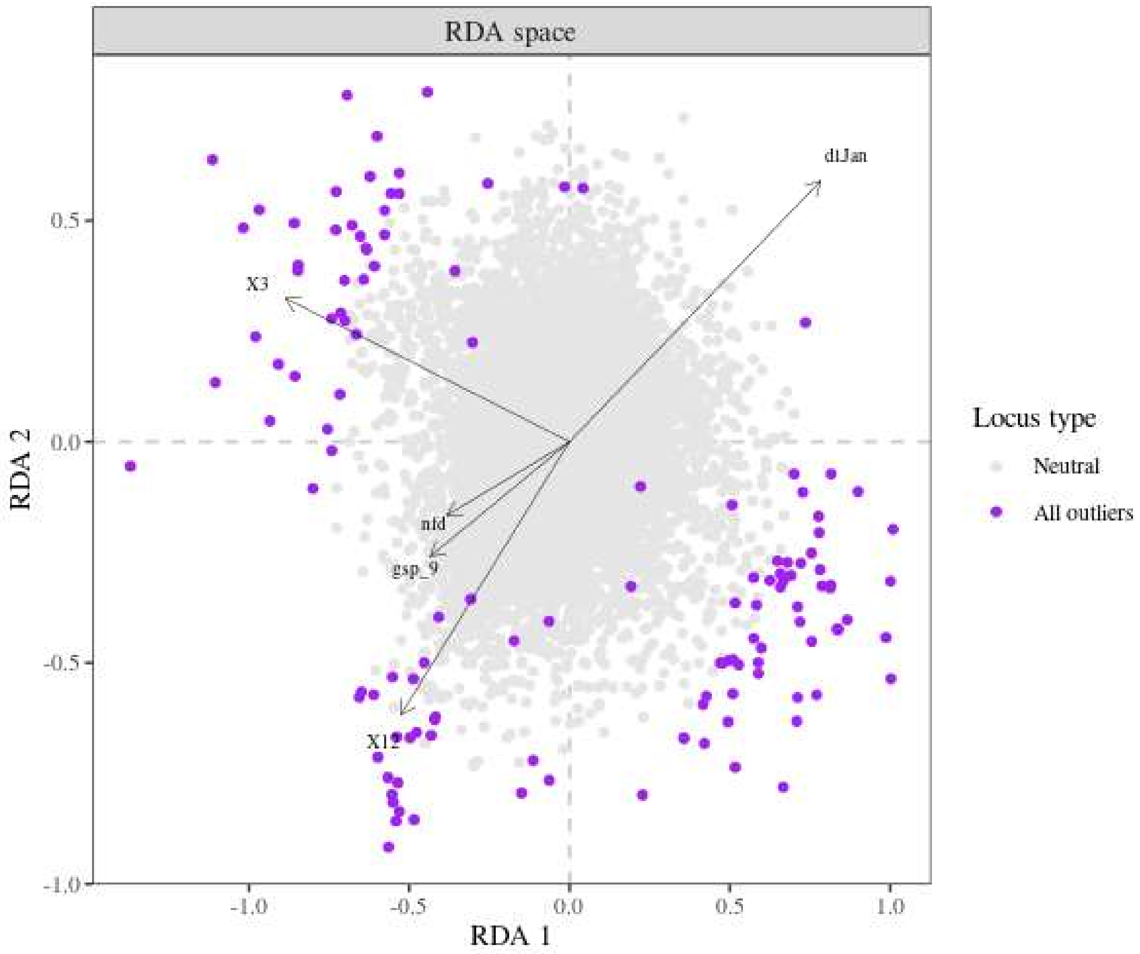


**Figure S4. RDA biplot showing outlier and neutral loci distribution in relation to chosen environmental variables on the first two axes.** Putatively neutral loci are shown in grey, while putatively adaptive loci are shown in purple.

**Table S1. All significant annotated outliers identified by at least one genome scan analysis.** Location within *Nothofagus pumilio* is indicated by contig and position. Annotation information was sourced from the UNIPROT database, including reference genome species, gene name (shorthand), gene function, and UniProt accession code. Colors of "Gene and function" column indicate gene function category, for which a legend is provided at the top of the chart. Synonymousness is indicated by S (synonymous) or NS (nonsynonymous). For each genome scan that identified this locus as a significant outlier, the qvalue is provided. For each outlier that BayPass identified, the Bayes Factor per environmental factor is provided when the value was significant (BF > 10).

| Gene Function Category | | |
|---|---|---|
| Stress response | Growth & Development | Ion transport & homeostasis |
| Synthesis-metabolism-photosynthesis | Transcriptional regulation | Other |

| *N. pumilio* location | | Annotation information | | | | | Significance values (qvalues) per analysis | | | | Bayes Factor (if >10) | | | |
|---|---|---|---|---|---|---|---|---|---|---|---|---|---|---|
| Contig | Pos | Ref genome species | Gene name | Gene and function | UniProt | Syn | pcadapt | LFMM2 | RDA | BayPass | GSP 9 | Ann Pre NFD | Isotherm | DLJ |
| chain_10152 | 1381 | Arabidopsis thaliana | PGIC | Sugar isomerase (SIS) family protein | P34795 | S | 1.18E-06 | 5.04E-08 | 2.66E-04 | 1.03E-07 | | | | 24.015 |
| chain_17554 | 602 | Carya illinoinensis | | myosin-J heavy chain-like | Q9C8T4 | S | 3.63E-08 | 8.96E-08 | 1.64E-04 | 3.64E-14 | | | 46.937 | 14.733 |
| chain_2392 | 1061 | Juglans regia | MYC2 | transcription factor MYC2 | Q39204 | S | 3.44E-08 | 1.07E-07 | 7.33E-04 | 2.86E-12 | 18.358 | | 19.852 | 12.109 |
| chain_4216 | 1080 | Arabidopsis thaliana | WRKY4 | WRKY DNA-binding protein 4 (WRKY4) | Q9XI90 | S | 2.63E-07 | 9.37E-04 | 1.83E-03 | 3.64E-13 | | | | 18.979 |
| chain_4216 | 1479 | Arabidopsis thaliana | WRKY4 | WRKY DNA-binding protein 4 (WRKY4) | Q9XI90 | S | 7.76E-07 | 5.30E-05 | 9.90E-04 | 7.14E-08 | | | 10.065 | 19.574 |
| chain_4370 | 996 | Arabidopsis thaliana | GLU1 | glutamate synthase 1 (GLU1) | Q9ZNZ7 | S | 1.19E-08 | 2.89E-04 | 3.03E-05 | 2.40E-09 | | | 20.043 | |
| chain_4914 | 201 | Arabidopsis thaliana | MYB12 | myb domain protein 12 (MYB12) | O22264 | S | 4.27E-10 | 2.97E-10 | 2.34E-05 | 0.00E+00 | | | 52.964 | |
| chain_5380 | 2377 | Juglans regia | | GUANYLATE-BINDING PROTEIN-LIKE 1 | F4I2I5 | S | 7.41E-05 | 1.58E-05 | 1.28E-03 | 1.58E-06 | | | | 25.896 |
| chain_707 | 585 | Arabidopsis thaliana | CYP73A5 | cinnamate-4-hydroxylase (C4H) | P92994 | S | 2.96E-03 | 1.20E-03 | 9.48E-03 | 5.99E-06 | | | | 24.522 |
| chain_747 | 927 | Arabidopsis thaliana | GLT1 | glutamate synthase 1 (GLU1, GLS1, GLUS, FD-GOGAT) | Q9LV03 | S | 1.19E-08 | 2.89E-04 | 3.03E-05 | 3.32E-09 | | | 25.204 | |
| chain_7686 | 1170 | Arabidopsis thaliana | DRB2 | dsRNA-binding protein 2 (DRB2) | Q9SKN2 | S | 3.93E-12 | 4.94E-04 | 4.30E-05 | 1.66E-06 | | | 14.202 | |
| NODE_13005 | 432 | Arabidopsis thaliana | T22E16.240 | Regulator of chromosome condensation (RCC1) family protein | Q9M2S1 | S | 3.47E-10 | 9.37E-04 | 1.42E-05 | 0.00E+00 | | | 20.391 | |
| NODE_206 | 1894 | Quercus lobata | ISA2 | isoamylase 2, chloroplastic | Q8L735 | S | 1.39E-04 | 4.83E-03 | 7.64E-03 | 1.38E-05 | | | 13.936 | |

| *N. pumilio* location | | Annotation information | | | | | Significance values (qvalues) per analysis | | | | Bayes Factor (if >10) | | | |
|---|---|---|---|---|---|---|---|---|---|---|---|---|---|---|
| Contig | Pos | Ref genome species | Gene name | Gene and function | UniProt | Syn | pcadapt | LFMM2 | RDA | BayPass | GSP 9 | Ann Pre NFD | Isotherm | DLJ |
| NODE_3033 | 288 | Arabidopsis thaliana | AAP6 | AMINO ACID PERMEASE 6 | P92934 | S | 3.66E-05 | 4.57E-06 | 2.13E-04 | 3.70E-08 | | | | 28.992 |
| NODE_76105 | 35 | Arabidopsis thaliana | CYP86A22 | cytochrome P450 86A22 | B3RFJ6 | S | 2.12E-20 | 6.97E-11 | 8.51E-07 | 0.00E+00 | 12.558 | | 18.412 | |
| NODE_7808 | 279 | Arabidopsis thaliana | | sugar transporter, putative (DUF1195) | Q8LFK6 | S | 5.85E-07 | 5.04E-08 | 2.18E-04 | 3.86E-09 | | | | 26.339 |
| chain_33088 | 826 | Juglans regia | FER3 | ferritin-3, chloroplastic | Q9LYN2 | NS | 6.75E-04 | 5.71E-07 | 5.32E-04 | 2.01E-11 | | | | 39.913 |
| chain_4895 | 2604 | Arabidopsis thaliana | GLR3.3 | glutamate receptor 3.3 (ATGLR3.3, GLR3.3) | Q9C8E7 | NS | 1.69E-18 | 1.66E-05 | 1.23E-06 | 8.18E-13 | | | 12.491 | |
| chain_5380 | 1917 | Arabidopsis thaliana | | GUANYLATE-BINDING PROTEIN-LIKE 1 | F4I2I5 | NS | 4.41E-04 | 1.03E-06 | 1.56E-03 | 8.82E-05 | | | 25.737 | |
| chain_6613 | 2642 | Quercus suber | NHX8 | sodium/hydrogen exchanger 8 | Q3YL57 | NS | 1.29E-09 | 5.30E-05 | 2.43E-05 | 2.63E-05 | | | 23.111 | |
| chain_7453 | 1118 | Quercus suber | PHS1 | alpha-1,4 glucan phosphorylase L-2 isozyme, chloroplastic/amyloplastic-like | Q9LIB2 | NS | 2.32E-07 | 1.61E-03 | 4.16E-03 | 0.00E+00 | | | | 15.481 |
| chain_9640 | 3371 | Arabidopsis thaliana | PPC4 | phosphoenolpyruvate carboxylase 4 (ATPPC4, PPC4) | Q8GVE8 | NS | 3.33E-04 | 1.62E-03 | 7.60E-03 | 8.51E-08 | | | 13.162 | |
| NODE_10100 | 253 | Arabidopsis thaliana | PUMP3 | mitochondrial uncoupling protein 3 (PUMP3/UCP3) | Q9XI74 | NS | 9.50E-15 | 3.26E-06 | 2.26E-06 | 0.00E+00 | | | 17.959 | |
| NODE_1604 | 1183 | Quercus suber | AKT1 | AKT1-like | Q38998 | NS | 2.00E-06 | 3.80E-06 | 7.72E-04 | 8.83E-11 | | | 18.874 | 37.845 |
| NODE_20727 | 497 | Arabidopsis thaliana | DXPS1 | 1-deoxy-D-xylulose 5-phosphate synthase 1 | F4IXL8 | NS | 3.28E-04 | 2.98E-04 | 7.45E-03 | 2.73E-08 | | | 22.702 | |
| NODE_256 | 1338 | Betula platyphylla | CYP74A | allene oxide synthase | Q96242 | NS | 1.09E-04 | 3.70E-03 | 1.29E-03 | 7.62E-05 | | | 26.254 | |
| NODE_4269 | 834 | Quercus lobata | CYCT1-3 | cyclin-T1-3-like | Q8LBC0 | NS | 2.99E-06 | 3.86E-04 | 1.88E-04 | 1.82E-06 | | | 10.65 | |
| NODE_50461 | 203 | Juglans regia | LRK10 | rust resistance kinase Lr10-like | P93604 | NS | 1.98E-03 | 1.11E-04 | 1.82E-03 | 4.76E-10 | | | 16.068 | |
| NODE_908 | 734 | Quercus suber | RH21 | DEAD-box ATP-dependent RNA helicase 21-like | P93008 | NS | 1.64E-14 | 2.24E-05 | 5.98E-07 | 0.00E+00 | | 11.777 | | |
| chain_1133 | 1296 | Arabidopsis thaliana | PER7 | Peroxidase superfamily protein | Q9SY33 | S | 5.31E-04 | 6.39E-03 | | 6.63E-08 | | | | 35.053 |
| chain_12200 | 1304 | Arabidopsis thaliana | CUV | RNA helicase family protein (EMB3011) | F4K2E9 | S | | 1.10E-04 | 8.85E-04 | 8.02E-07 | | | 18.394 | 15.57 |

| *N. pumilio* location | | Annotation information | | | | | Significance values (qvalues) per analysis | | | | Bayes Factor (if >10) | | | | |
|---|---|---|---|---|---|---|---|---|---|---|---|---|---|---|---|
| Contig | Pos | Ref genome species | Gene name | Gene and function | UniProt | Syn | pcadapt | LFMM2 | RDA | BayPass | GSP 9 | Ann Pre | NFD | Isotherm | DLJ |
| chain_16058 | 281 | Arabidopsis thaliana | CCD1 | nine-cis-epoxycarotenoid dioxygenase 3 (NCED3, ATNCED3, STO1, SIS7) | O65572 | S | 3.06E-10 | 1.68E-03 | 1.23E-04 | | | | | | |
| chain_1793 | 1302 | Arabidopsis thaliana | LOS1 | Ribosomal protein S5/Elongation factor G/III/V family protein (LOS1) | Q9ASR1 | S | 3.53E-04 | 1.36E-03 | | 1.04E-06 | | | 23.533 | | |
| chain_1793 | 1473 | Arabidopsis thaliana | LOS1 | Ribosomal protein S5/Elongation factor G/III/V family protein (LOS1) | Q9ASR1 | S | 5.50E-04 | 5.05E-03 | | 6.54E-03 | | | 11.963 | | |
| chain_18733 | 268 | Quercus suber | | glucan endo-1,3-beta-glucosidase 5 | Q9M088 | S | 4.18E-03 | | 6.09E-03 | 1.41E-05 | | 39.913 | | | |
| chain_2479 | 302 | Arabidopsis thaliana | | Hexokinase (ATHXK4, HKL2) | Q9LPS1 | S | 1.41E-09 | 4.91E-08 | | 0.00E+00 | | | 52.964 | | 14.903 |
| chain_45519 | 1123 | Arabidopsis thaliana | AP22.19 | Chaperone DnaJ-domain | F4JPR4 | S | 1.17E-09 | 5.01E-08 | | 0.00E+00 | 52.964 | 52.964 | | | |
| chain_52210 | 405 | Arabidopsis thaliana | RBOHH | laccase 17 (LAC17) | Q9FJD6 | S | 4.21E-05 | 9.99E-04 | | 1.88E-03 | 17.344 | | | | |
| chain_6538 | 1269 | Arabidopsis thaliana | EBF1 | EIN3-binding F box protein 1 (EBF1, FBL6) | Q9SKK0 | S | | 7.84E-04 | 6.25E-03 | 2.05E-05 | | | | | 27.662 |
| chain_8505 | 860 | Juglans microcarpa x Juglans regia | | phosphoglycerate mutase-like protein 4 | Q9SCS3 | S | 9.02E-03 | 7.38E-03 | | 1.27E-04 | | 14.46 | | 22.954 | |
| NODE_256 | 1654 | Betula platyphylla | CYP74A | allene oxide synthase | Q96242 | S | 2.45E-12 | 9.37E-04 | 4.05E-06 | | | | | | |
| NODE_3272 | 182 | Arabidopsis thaliana | CNGC4 | Cyclic nucleotide-gated ion channel 4 | Q94AS9 | S | 3.51E-03 | 6.76E-03 | 6.49E-03 | | | | | | |
| chain_11318 | 2942 | Quercus suber | CSLG3 | cellulose synthase-like protein G3 (LOC112016029) | Q0WVN5 | NS | 2.96E-06 | | 7.84E-04 | 5.39E-03 | | | | 13.039 | |
| chain_1175 | 3818 | Arabidopsis thaliana | MS2 | 5-methyltetrahydropteroyltriglutamate-homocysteine S-methyltransferase - like protein | Q9SRV5 | NS | 4.09E-03 | 1.59E-03 | | 1.51E-06 | | | | 19.085 | |
| chain_1793 | 798 | Arabidopsis thaliana | LOS1 | Ribosomal protein S5/Elongation factor G/III/V family protein | Q9ASR1 | NS | 1.23E-04 | 3.30E-03 | | 1.28E-03 | | | 18.25 | | |
| chain_24539 | 344 | Arabidopsis thaliana | BZR1 | Brassinosteroid signalling positive regulator (BZR1) family protein (BZR1) | Q8S307 | NS | 5.23E-04 | 1.73E-03 | | 3.32E-06 | | | | | 13.909 |

| *N. pumilio* location | | Annotation information | | | | | Significance values (qvalues) per analysis | | | | Bayes Factor (if >10) | | | | |
|---|---|---|---|---|---|---|---|---|---|---|---|---|---|---|---|
| Contig | Pos | Ref genome species | Gene name | Gene and function | UniProt | Syn | pcadapt | LFMM2 | RDA | BayPass | GSP 9 | Ann Pre | NFD | Isothern | DLJ |
| chain_5171 | 1021 | Arabidopsis thaliana | DL4875C | SBP (S-ribonuclease binding protein) family protein | Q8L7G9 | NS | | 8.06E-05 | 4.16E-03 | 1.49E-04 | | | | | 16.522 |
| chain_52210 | 236 | Arabidopsis thaliana | LAC17 | laccase 17 (LAC17) | Q9FJD5 | NS | 2.86E-07 | 1.58E-03 | 1.88E-04 | | | | | | |
| chain_560 | 460 | Quercus lobata | GDH3 | glycine cleavage system H protein 3, mitochondrial | Q9LQL0 | NS | 1.23E-04 | 2.84E-03 | 1.28E-03 | | | | | | |
| chain_6637 | 671 | Arabidopsis thaliana | LHCB2.2 | photosystem II light harvesting complex gene 2.2 (LHCB2.2, LHCB2) | Q9S7J7 | NS | 7.43E-03 | | 6.87E-03 | 1.51E-06 | | | | 20.881 | |
| chain_8494 | 399 | Arabidopsis thaliana | AGO4 | Argonaute family protein (AGO4, OCP11) | Q9ZVD5 | NS | | 1.23E-04 | 6.42E-03 | 2.87E-12 | | 52.964 | | | 34.764 |
| chain_8522 | 1657 | Arabidopsis thaliana | MED25 | phytochrome and flowering time regulatory protein (PFT1) (PFT1) | Q7XYY2 | NS | 6.70E-23 | 5.01E-08 | 2.90E-08 | | | | | | |
| chain_9104 | 3616 | Quercus lobata | BAM7 | beta-amylase 7 | O80831 | NS | 2.85E-03 | 8.46E-03 | | 1.19E-05 | | | | 15.548 | 16.742 |
| chain_9565 | 1507 | Juglans regia | BOB1 | protein BOBBER 1-like (chaperona / auxinas) | Q9LV09 | NS | 1.68E-03 | | 6.58E-03 | 2.38E-03 | | | | 10.267 | |
| NODE_11997 | 171 | Quercus suber | PYL4 | abscisic acid receptor PYL4-like | O80920 | NS | | 5.94E-05 | 3.62E-03 | 1.23E-06 | | | | | 32.391 |
| NODE_1583 | 1464 | Quercus robur | REF6 | lysine-specific demethylase REF6 | Q9STM3 | NS | 3.27E-04 | | 6.48E-03 | 3.16E-04 | | | | 19.869 | |
| NODE_1771C | 617 | Arabidopsis thaliana | ABH1 | ARM repeat superfamily protein (ENS, ABH1, CBP80, ATCBP80) | Q9SIU2 | NS | 9.11E-04 | 5.50E-03 | | 1.52E-12 | 10.217 | 11.89 | 11.623 | | |
| NODE_206 | 194 | Quercus lobata | ISA2 | isoamylase 2, chloroplastic | Q8L735 | NS | 1.56E-04 | | 9.49E-03 | 7.26E-06 | | | | 12.457 | 11.262 |
| NODE_2144 | 465 | Quercus suber | | probable protein phosphatase 2C 24 | Q9ZW21 | NS | 4.45E-10 | 2.39E-03 | | 6.82E-14 | 52.964 | 23.131 | | | |
| NODE_23626 | 36 | Prunus dulcis | KCS4 | 3-ketoacyl-CoA synthase 4 | Q9LN49 | NS | 4.90E-07 | 7.59E-03 | 2.42E-04 | | | | | | |
| NODE_8149 | 201 | Arabidopsis thaliana | MES1 | METHYL ESTERASE 1 | Q8S8S9 | NS | 3.44E-03 | 5.30E-05 | | 7.62E-05 | | | | | 12.25 |
| chain_1105 | 141 | Arabidopsis thaliana | SMT2 | 24-sterol C-methyltransferase (At1g20330) mRNA | Q39227 | S | 6.24E-07 | | 3.54E-03 | | | | | | |
| chain_1223 | 1117 | Quercus suber | ANTR6 | probable anion transporter 6, chloroplastic | Q3E9A0 | S | 2.90E-03 | | | 1.12E-05 | | | | 15.185 | |
| chain_12596 | 1753 | Quercus lobata | PHYA | phytochrome A | P14712 | S | | 3.91E-05 | | 3.89E-05 | | | | | 30.505 |

| *N. pumilio* location | | Annotation information | | | | | Significance values (qvalues) per analysis | | | | | Bayes Factor (if >10) | | | |
|---|---|---|---|---|---|---|---|---|---|---|---|---|---|---|---|
| Contig | Pos | Ref genome species | Gene name | Gene and function | UniProt | Syn | pcadapt | LFMM2 | RDA | BayPass | GSP 9 | Ann Pre | NFD | Isotherm | DLJ |
| chain_13219 | 428 | Arabidopsis thaliana | MKK3 | mitogen-activated protein kinase kinase 3 | O80396 | S | 5.66E-07 | | 7.84E-04 | | | | | | |
| chain_1383 | 432 | Arabidopsis thaliana | GGR | geranylgeranyl reductase (GGR) | Q39108 | S | | | 3.13E-03 | 8.54E-08 | | | 15.459 | 23.37 | 10.742 |
| chain_1719 | 912 | Quercus suber | ZFN3 | zinc finger CCCH domain-containing protein ZFN-like | Q8L7N8 | S | | 3.46E-03 | | 5.71E-05 | | | | | 36.049 |
| chain_18152 | 230 | Quercus lobata | CTL2 | chitinase 2-like | Q9LSP9 | S | | 7.26E-04 | | 8.09E-03 | | | | | 13.743 |
| chain_1867 | 481 | Quercus robur | APL3 | glucose-1-phosphate adenylyltransferase large subunit 3 | P55231 | S | | 5.61E-04 | | 3.89E-05 | | | 10.876 | | 24.522 |
| chain_3018 | 720 | Juglans microcarpa x Juglans regia | BCA2 | carbonic anhydrase 2-like | P42737 | S | | 6.61E-04 | | 7.20E-05 | | | | 23.17 | |
| chain_37834 | 3636 | Juglans regia | BFRUCT3 | acid beta-fructofuranosidase-like | Q43348 | S | 6.43E-03 | 9.63E-03 | | | | | | | |
| chain_5883 | 2781 | Quercus lobata | ISA3 | isoamylase 3, chloroplastic | Q9M0S5 | S | 2.66E-04 | | 6.17E-03 | | | | | | |
| chain_6657 | 1989 | Arabidopsis thaliana | HAB1 | homology to ABI1 (HAB1) | Q9CAJ0 | S | 1.35E-06 | | 1.92E-03 | | | | | | |
| chain_7367 | 914 | Carya illinoinensis | TPS9 | alpha,alpha-trehalose-phosphate synthase [UDP-forming] 9 | Q9LRA7 | S | | 2.39E-03 | | 1.76E-04 | | | | | 17.552 |
| chain_7982 | 608 | Arabidopsis thaliana | NAC082 | NAC domain containing protein 82 | Q9FY82 | S | | 5.13E-04 | | 7.62E-05 | | | | | 17.495 |
| chain_8745 | 1180 | Quercus lobata | ACO4 | 1-aminocyclopropane-1-carboxylate oxidase-like | Q06588 | S | 6.29E-05 | | 5.27E-03 | | | | | | |
| NODE_1583 | 154 | Quercus robur | REF6 | lysine-specific demethylase REF6 | Q9STM3 | S | 7.49E-07 | | 5.12E-04 | | | | | | |
| NODE_1822 | 1520 | Quercus suber | | Nucleolin-like | Q8LFA8 | S | | 8.46E-03 | | 6.58E-07 | | | | | 22.249 |
| NODE_23626 | 94 | Prunus dulcis | KCS4 | 3-ketoacyl-CoA synthase 4 | Q9LN49 | S | 2.15E-07 | | 3.13E-03 | | | | | | |
| NODE_2863 | 329 | Arabidopsis thaliana | SCPL8 | SNG1 / SCPL8 | Q8RUW5 | S | | 8.44E-04 | | 1.12E-04 | | | | | 29.796 |
| NODE_3134 | 1038 | Juglans microcarpa x Juglans regia | SS4 | probable starch synthase 4, chloroplastic/amyloplastic | Q0WVX5 | S | 1.05E-04 | | 7.06E-03 | | | | | | |

| *N. pumilio* location | | Annotation information | | | | | Significance values (qvalues) per analysis | | | | | Bayes Factor (if >10) | | |
|---|---|---|---|---|---|---|---|---|---|---|---|---|---|---|
| Contig | Pos | Ref genome species | Gene name | Gene and function | UniProt | Syn | pcadapt | LFMM2 | RDA | BayPass | GSP 9 | Ann Pre NFD | Isotherm | DLJ |
| NODE_4866 | 798 | Quercus suber | | cytochrome P450 71A4-like | A0A654F | S | 1.02E-04 | | 2.93E-03 | | | | | |
| NODE_51 | 766 | Arabidopsis thaliana | CDF3 | cycling DOF factor 3 (CDF3) | Q8LFV3 | S | 1.31E-03 | | | 3.91E-07 | | | 10.742 | |
| chain_10180 | 583 | Quercus robur | FZL | probable transmembrane GTPase FZO-like, chloroplastic (LOC126710593), mRNA | Q1KPV0 | NS | 2.70E-03 | | | 8.56E-05 | | | 10.787 | |
| chain_11318 | 2426 | Quercus suber | CSLG3 | cellulose synthase-like protein G3 (LOC112016029) | Q0WVN5 | NS | 2.83E-04 | | 9.97E-03 | | | | | |
| chain_11868 | 88 | Arabidopsis thaliana | SRFR1 | Tetratricopeptide repeat (TPR)-like superfamily protein (SRFR1) | F4JS25 | NS | 7.91E-07 | | 1.92E-04 | | | | | |
| chain_12349 | 734 | Quercus lobata | FUT1 | galactoside 2-alpha-L-fucosyltransferase-like | Q9SWH5 | NS | | 2.40E-03 | | 1.72E-03 | | | | 15.526 |
| chain_13578 | 435 | Quercus suber | ARF4 | auxin response factor 4 | Q9ZTX9 | NS | 2.40E-07 | | 1.57E-03 | | | | | |
| chain_15177 | 1141 | Quercus suber | NLP7 | protein NLP7-like | Q84TH9 | NS | 2.36E-04 | | 4.38E-03 | | | | | |
| chain_15308 | 358 | Arabidopsis thaliana | BGLU40 | beta glucosidase 40 (BGLU40) | Q9FZE0 | NS | 7.19E-04 | | 2.30E-03 | | | | | |
| chain_18733 | 616 | Quercus suber | | glucan endo-1,3-beta-glucosidase 5 | Q9M088 | NS | 8.50E-03 | | 7.50E-03 | | | | | |
| chain_2451 | 418 | Quercus lobata | RBCX2 | chaperonin-like RbcX protein 2, chloroplastic | Q8L9X2 | NS | 6.11E-04 | | | 6.92E-08 | | | 28.136 | |
| chain_2761 | 1325 | Juglans regia | VIL2 | VIN3-like protein | Q9SUM4 | NS | 1.34E-05 | | 2.64E-03 | | | | | |
| chain_2796 | 781 | Quercus lobata | AKR4C9 | NADPH-dependent aldo-keto reductase, chloroplastic-like | Q0PGJ6 | NS | | 9.08E-04 | 2.78E-03 | | | | | |
| chain_30268 | 1162 | Quercus robur | UVR8 | ultraviolet-B receptor UVR8 | Q9FN03 | NS | 1.23E-04 | | 5.59E-03 | | | | | |
| chain_3519 | 409 | Quercus robur | PFD1 | prefoldin subunit 1 | Q94AF7 | NS | 5.30E-03 | | 2.89E-03 | | | | | |
| chain_39074 | 2778 | Quercus lobata | LARP1A | la-related protein 1A | Q940X9 | NS | 4.07E-05 | | 3.13E-03 | | | | | |
| chain_4895 | 1942 | Arabidopsis thaliana | GLR3.3 | glutamate receptor 3.3 (ATGLR3.3, GLR3.3) | Q9C8E7 | NS | 1.58E-05 | | 1.89E-03 | | | | | |
| chain_5380 | 104 | Juglans regia | | GUANYLATE-BINDING PROTEIN-LIKE 1 | F4I2I5 | NS | 1.98E-03 | 6.93E-03 | | | | | | |
| chain_5907 | 3557 | Quercus robur | FTIP3 | FT-interacting protein 3 | Q9M2R0 | NS | 8.17E-03 | | | 1.36E-04 | | | 24.522 | |

| *N. pumilio* location | | Annotation information | | | | | Significance values (qvalues) per analysis | | | | Bayes Factor (if >10) | | | | |
|---|---|---|---|---|---|---|---|---|---|---|---|---|---|---|---|
| Contig | Pos | Ref genome species | Gene name | Gene and function | UniProt | Syn | pcadapt | LFMM2 | RDA | BayPass | GSP 9 | Ann Pre | NFD | Isotherm | DLJ |
| chain_5997 | 1397 | Arabidopsis thaliana | AHK3 | histidine kinase 3 (AHK3, HK3) | Q9C5U1 | NS | 6.67E-06 | | 5.50E-03 | | | | | | |
| chain_6613 | 2770 | Quercus suber | NHX8 | sodium/hydrogen exchanger 8 | Q3YL57 | NS | 1.56E-03 | | | 8.13E-06 | | | | 16.762 | |
| chain_7063 | 476 | Juglans microcarpa x Juglans regia | COG0212 | 5-formyltetrahydrofolate cyclo-ligase-like protein COG0212 | Q9SRE0 | NS | 1.13E-04 | | 4.24E-03 | | | | | | |
| chain_8309 | 2487 | Juglans regia | CAMTA2 | calmodulin-binding transcription activator 2-like | Q6NPP4 | NS | 7.18E-06 | | 3.76E-03 | | | | | | |
| NODE_1219 | 1506 | Quercus suber | PEPKR2 | serine/threonine-protein kinase PEPKR2 (catalytic activity) | Q8W490 | NS | | 1.42E-03 | | 3.74E-03 | | | | | 12.524 |
| NODE_12254 | 250 | Carya illinoinensis | MSSP2 | monosaccharide-sensing protein 2 | Q8LPQ8 | NS | 2.19E-03 | | | 1.28E-06 | | | | 22.101 | |
| NODE_12254 | 683 | Carya illinoinensis | MSSP2 | monosaccharide-sensing protein 2 | Q8LPQ8 | NS | 1.93E-05 | | 7.17E-03 | | | | | | |
| NODE_1682 | 1727 | Quercus suber | WRKY33 | WRKY transcription factor 33 | Q8S8P5 | NS | 1.99E-03 | | | 2.67E-08 | | | | | 17.382 |
| NODE_20566 | 106 | Quercus robur | | probable sarcosine oxidase | Q9SJA7 | NS | 9.81E-08 | | 6.27E-03 | | | | | | |
| NODE_2062 | 1626 | Juglans microcarpa x Juglans regia | VIP2 | protein PAF1 homolog | F4HQA1 | NS | 9.68E-05 | | 7.61E-03 | | | | | | |
| NODE_2144 | 402 | Quercus suber | | probable protein phosphatase 2C 24 | Q9ZW21 | NS | 1.93E-05 | | | 1.28E-06 | 18.927 | 15.681 | 12.491 | | |
| NODE_2144 | 444 | Quercus suber | | probable protein phosphatase 2C 24 | Q9ZW21 | NS | 6.67E-06 | | | 1.50E-05 | | | 38.76 | | |
| NODE_2515 | 1000 | Quercus lobata | GT14 | probable xyloglucan galactosyltransferase GT14 (xyloglucan biosynthesis) | Q84R16 | NS | | 3.13E-03 | | 8.80E-05 | 16.047 | 20.391 | 15.919 | | |
| NODE_3734 | 435 | Arabidopsis thaliana | DURF2 | E3 ubiquitin-protein ligase RDUF2 (ABA mediated stress responses) | Q940T5 | NS | 1.17E-03 | | | 3.35E-10 | | | 43.918 | | |
| NODE_4037 | 556 | Quercus robur | | AAA-ATPase At3g50940-like (cathalytic activity) | Q147F9 | NS | 4.94E-03 | | 5.03E-03 | | | | | | |
| NODE_50461 | 134 | Juglans regia | LRK10 | rust resistance kinase Lr10-like | P93604 | NS | | | 4.69E-03 | 7.62E-05 | | 14.585 | | | |
| NODE_5764 | 71 | Quercus suber | CYP71A24 | cytochrome P450 71A24-like | Q9STK9 | NS | | 5.30E-05 | | 4.54E-04 | | | | 20.723 | |
